# Autophagy induction requires the suppression of potassium influx mediated by phosphatases

**DOI:** 10.64898/2026.02.27.707896

**Authors:** Natsu Matsumoto, Suzuna Akema, Shigehisa Akiyama, Yasuhiro Araki, Takeshi Noda

## Abstract

Potassium is an essential element for all living organisms. As the most abundant intracellular cation, it is vital for osmoregulation, maintenance of membrane potential, macromolecule transport and enzyme function. In this study, we identify potassium homeostasis as a previously unknown regulator of autophagy through the phosphatase activities of Ppz1 and Ppz2. We find that overexpression of either Ppz1 or Ppz2 triggers autophagy under nutrient-replete conditions, whereas the loss of both causes severe autophagy defects. Strikingly, deletion of the potassium transporters Trk1 and Trk2, which are substrates of Ppz1 and Ppz2-mediated dephosphorylation, restores autophagic activity in *ppz1*Δ *ppz2*Δ cells. Furthermore, intracellular potassium concentrations declined during autophagy induction in wild-type cells but remained stable in *ppz1*Δ *ppz2*Δ mutants. Collectively, these findings establish Ppz1 and Ppz2 as pivotal regulators of autophagy and underscore intracellular potassium reduction as a primary determinant of this process.

**Summary statement:** This study demonstrates that potassium regulation by the phosphatases, Ppz1 and Ppz2, which suppress the major potassium transporters Trk1 and Trk2, is indispensable for autophagy induction in yeast.

## Introduction

Macroautophagy, hereafter called autophagy, is a highly conserved eukaryotic intracellular degradation pathway conserved from yeast to humans. By facilitating the rapid breakdown of large quantities of cytosolic material including organelles, autophagy allows for the recycling of cellular materials, supporting survival during starvation, can clear misfolded or aggregated proteins, and is even able to destroy invading pathogens (Klionsky et al., 2016; Mizushima and Komatsu, 2011) Substrates are engulfed by an expanding double-membrane structure known as the isolation membrane (or phagophore), which subsequently grows into a cup-shaped membranous structure that ultimately closes to yield a completed autophagosome (Ohsumi, 2014). The autophagosome then fuses with the vacuolar membrane, releasing its cargo for degradation by hydrolases within the vacuolar lumen.

The biogenesis of the autophagosome is a complex, multi-step process orchestrated by a conserved machinery of at least 19 core *ATG* gene products (Kotani et al., 2023). Accumulating evidence indicates that the fine-tuned control of autophagy is coordinated through phosphorylation. While the roles of protein kinases like TORC1 and Atg1 in autophagy are well established (Licheva et al., 2022), little is known about protein phosphatases, which antagonize kinase activity through reversible dephosphorylation. Recent studies in yeast have identified several phosphatases that regulate the autophagic machinery. Specifically, Ptc2, Ptc3, and Cdc14 have been found to dephosphorylate the Atg1 complex to promote autophagy (Feng et al., 2022; Memisoglu et al., 2019). PP2A has also been reported to regulate autophagy, although this is controversial as studies have reported conflicting results (Yeasmin et al., 2016; Yorimitsu et al., 2009). Besides the Atg1 complex, numerous Atg proteins are phosphorylated (Leutert et al., 2023); however, the precise impact of these post-translational modifications on autophagy remains unclear as the specific kinases and phosphatases responsible remain poorly characterized.

Beyond protein-based regulation, ion homeostasis has emerged as another modulator of autophagy. In both yeast and mammalian cells, fluctuations in the concentrations of intracellular ions, most notably potassium (K^+^) ions, as well as chloride (Cl^-^) and lysosomal/vacuolar pH, have been shown to modulate autophagic activity cascades (Hosogi et al., 2014; Rangarajan et al., 2020; Williams et al., 2008). However, how ions are involved in the regulation of autophagy at a mechanistic level is poorly understood, both in terms of the role of specific ionic species, as well as their availability and concentration within the cell.

In this study, we report a gain-of-function screen for phosphatases which identifies the paralogs Ppz1 and Ppz2 as capable of inducing autophagy upon overexpression. Notably, we further find that the simultaneous deletion of both *PPZ1* and *PPZ2* genes results in a near-complete block of autophagy, demonstrating for the first time the involvement of Ppz1 and Ppz2 in the regulation of autophagy. Mechanistic analyses indicate that Ppz1 and Ppz2 promote autophagy initiation by facilitating Atg1 kinase activity. We provide evidence that Ppz1 and Ppz2 function downstream of TORC1: overexpression of Ppz1 or Ppz2 induced autophagy under nutrient-rich conditions without detectable changes in TORC1 activity, and *ppz1*Δ *ppz2*Δ cells exhibited persistent autophagic defects even when TORC1 was inhibited by rapamycin. Because Ppz1 and Ppz2 are known regulators of K⁺ transporter Trk1 and Trk2, we also considered a potential role for potassium homeostasis. Genetic and physiological manipulations of K⁺ transport further demonstrate the importance of potassium availability in this pathway. Together, these results indicate that Ppz1 and Ppz2-dependent suppression of K⁺ influx is required for efficient Atg1 activation and autophagy initiation.

## Results

### Overexpression of either Ppz1 or Ppz2 initiates autophagy independently of TORC1 inactivation

In order to systematically screen for phosphatases that modulate autophagy, we generated a library of yeast strains overexpressing phosphatases and assessed autophagic flux under nutrient-replete conditions. *Saccharomyces cerevisiae* encodes 43 protein phosphatases (Offley and Schmidt, 2019) with a range of substrate specificities; of these, we excluded non-protein-targeting and atypical enzymes (Ppn2, Ymr1, Rtr1 and Rtr2), leaving us with a library of 39 phosphatase overexpression strains. Overexpression was achieved using the hormone-based gene expression Z3EV/Z4EV system (McIsaac et al., 2013), allowing conditional expression of a protein of interest upon supplementation of the physiologically inert hormone β-estradiol. Following treatment of cells with β-estradiol for 3 h, autophagy activity was determined using the GFP-Atg8 cleavage assay (Klionsky et al., 2007). Using this assay, the induction of autophagy is easily detected as a free GFP band appears in western blots, as observed when cells are treated with rapamycin, which acts by inhibiting TORC1 kinase activity and thereby strongly induces autophagy (Fig. 1A).

**Figure 1.**
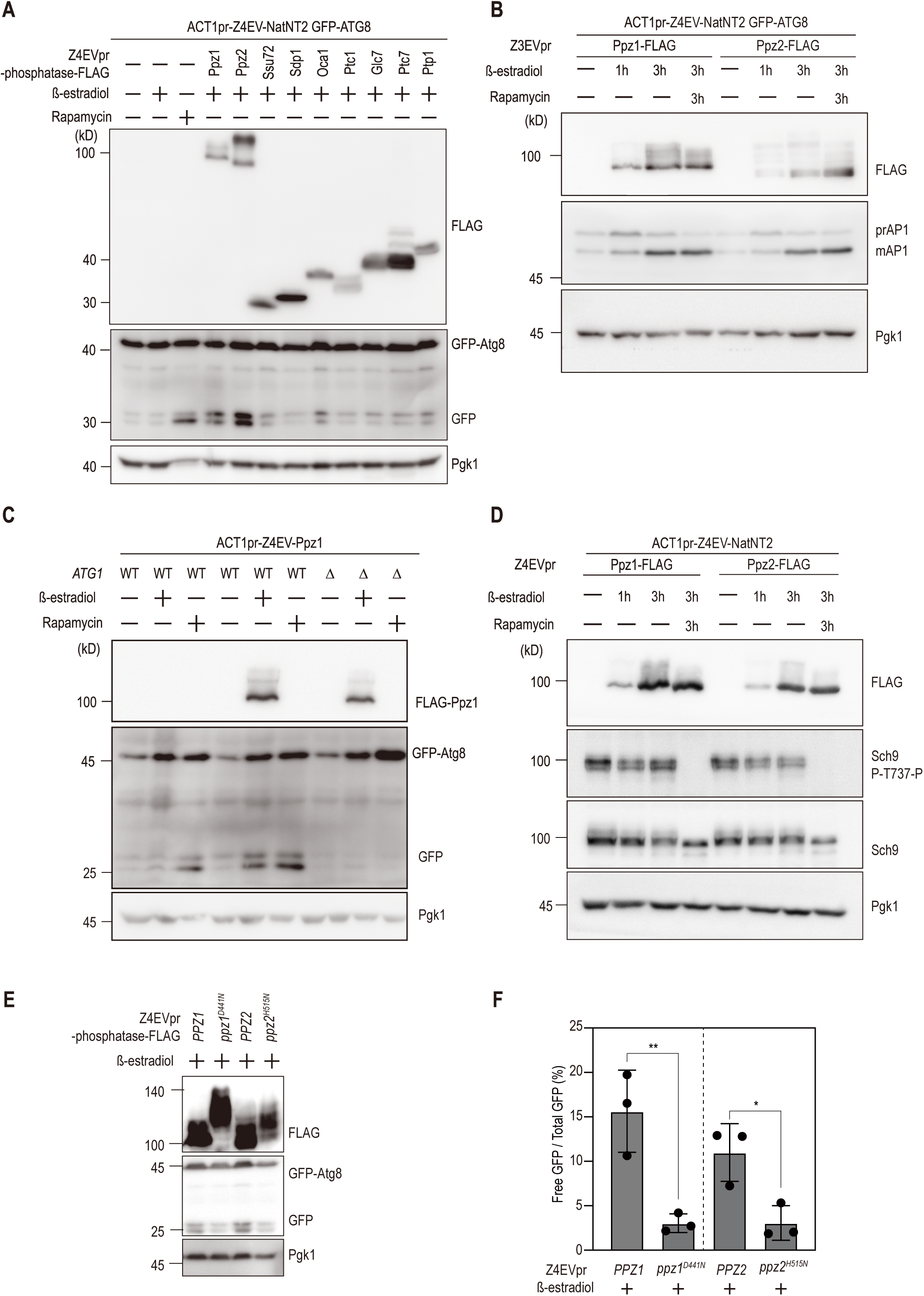
Overexpression of either Ppz1 or Ppz2 induces autophagy without inactivation of TORC1. (A) Cells were treated with 2 µg/mL β-estradiol or 0.2 µg/mL rapamycin for 3 h. Lysates were analyzed by immunoblotting with anti-FLAG, anti-GFP, and anti-Pgk1 antibodies. Each phosphatase was fused to a FLAG tag to confirm overexpression. Rapamycin served as a positive control. (B) Wild-type cells and cells overexpressing Ppz1 or Ppz2 were collected after 3 h of treatment with β-estradiol, rapamycin, or both. Lysates were analyzed by immunoblotting with anti-FLAG, anti-Ape1, and anti-Pgk1 antibodies. (C) Wild-type and *atg1*Δ cells, both overexpressing Ppz1, were harvested following 3 h of treatment with either β-estradiol or rapamycin. Lysates were analyzed by immunoblotting with anti-FLAG, anti-GFP, and anti-Pgk1 antibodies. (D) Cells overexpressing either Ppz1 or Ppz2 were collected after 3 h of treatment with β-estradiol or rapamycin and analyzed by immunoblotting with anti-FLAG, anti-Sch9-phosphoT737, Sch9 and anti-Pgk1 antibodies. (E) Cells overexpressing either Ppz1 or Ppz2 and their respective catalytically inactive mutants were collected after 3 h of treatment with β-estradiol and analyzed by immunoblotting with anti-FLAG, anti-GFP, and anti-Pgk1 antibodies. (F) Quantification of Fig. 1E. The ratio of free GFP to total GFP signal intensity was used to compare autophagy flux. Data are presented as mean values ± s.d. from three independent experiments. Unpaired t-test: \*\**P* < 0.001, \**P* < 0.05

This screen identified two candidates which markedly increased autophagy activity: Ppz1 and Ppz2 (Fig. 1A; see Table S1 for complete screening results). Ppz1 and Ppz2 are paralogs which functionally redundant (Posas et al., 1993). To independently corroborate the induction of autophagy, we monitored the proteolytic processing of aminopeptidase I (Ape1). Ape1 is a well-established cargo that is delivered to the vacuole and matured via both the cytoplasm-to-vacuole targeting (Cvt) pathway and autophagy. While cytosolic precursor (prApe1) is constitutively transported via the Cvt pathway, its trafficking is significantly augmented by autophagy, leading to the efficient accumulation of the mature enzyme (mApe1) (Baba et al., 1997; Scott et al., 1996). This analysis also demonstrates that the overexpression of Ppz1 or Ppz2 strongly induces autophagy, as indicated by Ape1 maturation (Fig. 1B). To confirm that these results reflect the activity of the canonical autophagy pathway, we assessed GFP-Atg8 cleavage following the deletion of *ATG1*, the key kinase that is necessary for autophagy initiation (Matsuura et al., 1997). Deletion of *ATG1* completely abolished GFP-Atg8 cleavage, demonstrating that the cleavage induced by Ppz1 overexpression is strictly autophagy-dependent (Fig. 1C).

To determine whether the induction of autophagy by Ppz1 or Ppz2 is independent of TORC1 inactivation, we first assessed the phosphorylation status of canonical TORC1 substrates. Under nutrient-replete conditions, TORC1 directly phosphorylates Sch9 at Thr737; this modification is abolished upon TORC1 inhibition (Urban et al., 2007). Notably, Sch9 phosphorylation remained unaffected by the overexpression of Ppz1 or Ppz2 (Fig. 1D). Similarly, Atg13, which plays a critical role in autophagy induction when dephosphorylated (Kamada et al., 2010), showed no change in phosphorylation upon Ppz1 or Ppz2 induction (Supplemental Fig. 1A). These results suggest that Ppz1 and Ppz2 (hereafter Ppz1/2) neither function as upstream antagonists of TORC1 activity nor act as direct phosphatases mediating Atg13 dephosphorylation. Furthermore, while the overexpression of wild-type Ppz1 or Ppz2 robustly induced GFP-Atg8 cleavage, this effect was completely abrogated when catalytically inactive mutations were overexpressed under the same conditions (Fig. 1E, F). Collectively, these findings demonstrate that Ppz1/2 overexpression triggers autophagy in a phosphatase activity-dependent manner that operates independently of the canonical TORC1-Atg13 axis.

The catalytically inactive mutants of Ppz1 and Ppz2 exhibited reduced electrophoretic mobility compared to their wild-type counterparts (Fig. 1E). We speculated that this mobility shift may result from hyperphosphorylation of these proteins. To test this, we treated Ppz1 and its mutant with λ-phosphatase *in vitro*. Upon treatment, the higher-molecular-weight band of the inactive mutant collapsed to a position corresponding to that of the wild-type protein (Supplemental Fig. 1B). These results indicate that the catalytically inactive Ppz1 is hyperphosphorylated, suggesting that Ppz1/2 may exert autodephosphorylation or mutual dephosphorylation activities. These findings imply a potential self-regulatory mechanism, whereby constitutively phosphorylation of Ppz1 and Ppz2 under nutrient-rich conditions is counteracted by the phosphatase activity of these proteins.

### *ppz1*Δ *ppz2*Δ cells exhibit a defect in autophagy upon TORC1 inactivation

To further examine the function of Ppz1 and Ppz2 in autophagy, we next constructed deletion strains lacking the encoding genes. Autophagic flux was quantified using the alkaline phosphatase (ALP) assay, which monitors the vacuolar delivery and subsequent hydrolytic activation of the truncated Pho8Δ60 reporter (Noda and Ohsumi, 1998). In wild-type cells, ALP activity increased robustly in response to nitrogen starvation, whereas this increase was completely abolished in the autophagy-deficient *atg2*Δ strain (Fig. 2A), reflecting the lack of autophagy in these cells. While both the *ppz1*Δ and *ppz2*Δ single mutants showed similar ALP activity to WT cells, the *ppz1*Δ *ppz2*Δ double mutant displayed a profound defect, retaining only approximately 20% of the starvation-induced ALP activity in comparison to wild-type cells (Fig. 2A). In ALP assay, Ppz1 and Ppz2 function redundantly. We also performed the GFP-Atg8 cleavage assay to assess the effect of these deletions. Cells lacking *PPZ1* exhibited a marked reduction in GFP-Atg8 cleavage (roughly 50% of wild-type levels), whereas cleavage in *ppz2*Δ cells remained largely unaffected. Notably, in the GFP-Atg8 cleavage assay, the *ppz1*Δ *ppz2*Δ double mutant exhibited a phenotype similar to that of the *ppz1*Δ single mutant, suggesting that Ppz1 is the dominant contributor in this context (Fig. 2B, 2C). Importantly, plasmid-borne expression of the wild-type *PPZ1* gene completely rescued the autophagic defect in the double mutant strain (Fig. 2D, 2E), confirming that the observed phenotype is specifically attributable to the loss of Ppz1/2 activity.

**Figure 2.**
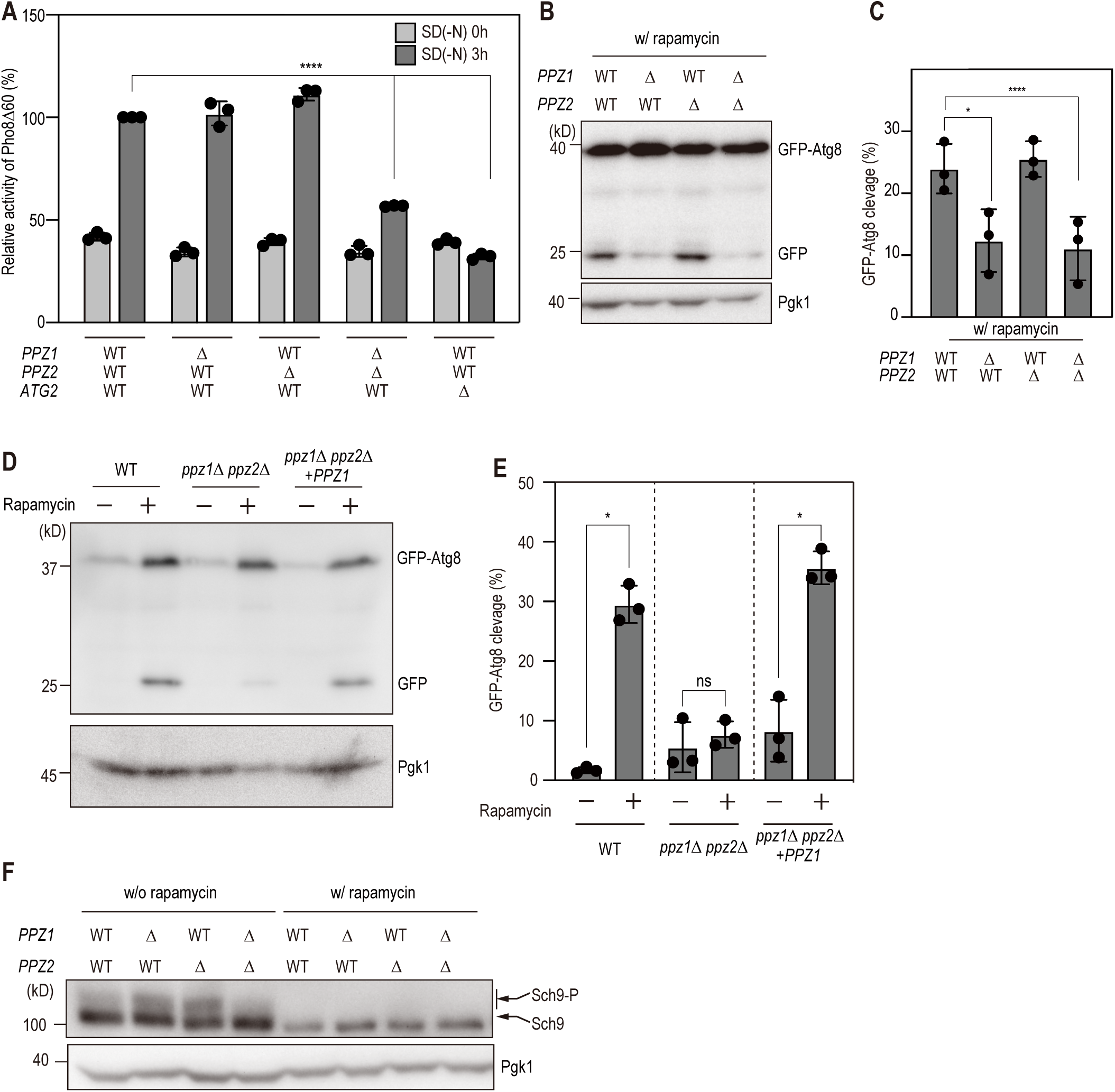
Loss of PPZ1 and PPZ2 impairs autophagy despite TORC1 inhibition. (A) Cells were incubated in YPD or SD-N medium for 3 h and subjected to an ALP assay at the indicated time points (n = 3). The y-axis shows relative ALP activity normalized to wild type (WT) treated with rapamycin (=100). Tukey’s multiple comparison test: \*\*\*\**P* < 0.0001 (B) Wild-type cells and cells lacking *PPZ1*, *PPZ2*, or both were collected after 3 h of treatment with 0.2 µg/mL rapamycin. Lysates were analyzed by immunoblotting with anti-GFP and anti-Pgk1 antibodies. (C) Quantification of Fig. 2B. The ratio of free GFP to total GFP was quantified in each strain to assess autophagy flux. Data are presented as mean values ± s.d. from three independent experiments. Tukey’s multiple comparison test: \*\*\*\**P* < 0.0001, \**P* < 0.005. (D) Wild-type cells, *ppz1*Δ *ppz2*Δ cells, and *ppz1*Δ *ppz2*Δ cells complemented with *PPZ1* were collected after 3 h of treatment with 0.2 µg/mL rapamycin. Lysates were analyzed by immunoblotting with anti-FLAG, anti-GFP, and anti-Pgk1 antibodies. (E) Quantification of Fig. 2D. The ratio of free GFP to total GFP was quantified in each strain to assess autophagy flux. Data are presented as mean values ± s.d. from three independent experiments. Paired two-tailed t-test with Holm–Šidák correction: *P* < 0.05; ns, *P* ≥ 0.05. (F) Wild-type cells and cells lacking *ppz1*, *ppz2*, or both were collected after 3 h of treatment with 0.2 µg/mL rapamycin. Lysates were analyzed by immunoblotting with anti-Sch9 and anti-Pgk1 antibodies.

We next examined the cellular localization of two autophagic markers, Atg1 and Atg8, to gain further insight into the Ppz1/2-mediated autophagy regulation. Under nutrient starvation conditions, both GFP-tagged Atg1 and Atg8 are typically sequestered into autophagosomes and delivered to the vacuole, where cleavage of the GFP moiety results in diffuse fluorescence within the vacuolar lumen (Kirisako et al., 2000; Kraft et al., 2012). However, microscopic analysis of the *ppz1*Δ *ppz2*Δ strain revealed a complete lack of intravacuolar GFP signal even after 3 hours of rapamycin treatment (Supplemental Fig. 2A, B). These findings demonstrate that the loss of Ppz1/2 results in a severe blockade in the autophagic trafficking of these essential proteins to the vacuole. Notably, despite the severe autophagy defect, the TORC1 substrate Sch9 was normally dephosphorylated in *ppz1*Δ *ppz2*Δ cells upon rapamycin treatment, confirming that TORC1 signaling is effectively suppressed even in the absence of these phosphatases (Fig. 2F). Collectively, these findings demonstrate that the functionally redundant phosphatases Ppz1 and Ppz2 are crucial for the induction of autophagy, functioning downstream of or independently from TORC1. This suggests that Ppz1 and Ppz2 are likely required for a key step in the autophagy mechanism, such as autophagosome biogenesis or autophagosome-vacuole fusion.

### Ppz1 and Ppz2 are required for Vps34 phosphorylation by Atg1

Following the induction of autophagy, the majority of core Atg proteins assemble at the pre-autophagosomal structure (PAS), a perivacuolar domain from which the autophagosome is formed (Suzuki et al., 2001; Suzuki et al., 2007). The severe impairment of autophagy in the *ppz1*Δ *ppz2*Δ mutant led us to investigate whether the hierarchical recruitment of core Atg proteins to the PAS was compromised. We constructed yeast strains in which the endogenous *ATG1* and *ATG2* genes were C-terminally tagged with GFP at their native chromosomal loci. Fluorescence microscopy revealed that the subcellular localization of both proteins was indistinguishable between wild-type and *ppz1*Δ *ppz2*Δ strains, as both proteins consistently assembled into characteristic puncta representing the PAS (Fig. 3A-D). However, as the frequency of Atg2-GFP puncta was marginally elevated in *ppz1*Δ *ppz2*Δ cells, we precisely quantified the intensity of the Atg2-GFP signal at the PAS (defined as Atg17-2×mCherry-positive puncta) (Suzuki et al., 2007). We found that the Atg2-GFP intensity in the *ppz1*Δ *ppz2*Δ strain was slightly higher than that in wild-type cells, although this increase was not significant (Supplemental Fig. 3A, B). As Atg18, an Atg2-binding partner, accumulates aberrantly at the PAS in mutants deficient in Atg1-mediated phosphorylation of Vps34, the catalytic subunit of Phosphatidylinositol 3-kinase complex (Lee et al., 2023), we next assessed whether the loss of Ppz1/2 impacts Vps34 phosphorylation. Western blot analysis revealed that Vps34 phosphorylation was markedly diminished in *ppz1*Δ *ppz2*Δ cells compared with wild-type cells (Fig. 3E, F). Since Atg1 has been identified as the direct kinase for Vps34 (Lee et al., 2023), our data suggest that the loss of Ppz1/2 is correlated with a profound impairment of Atg1 kinase activity, thereby disrupting the downstream signaling required for autophagosome formation.

**Figure 3.**
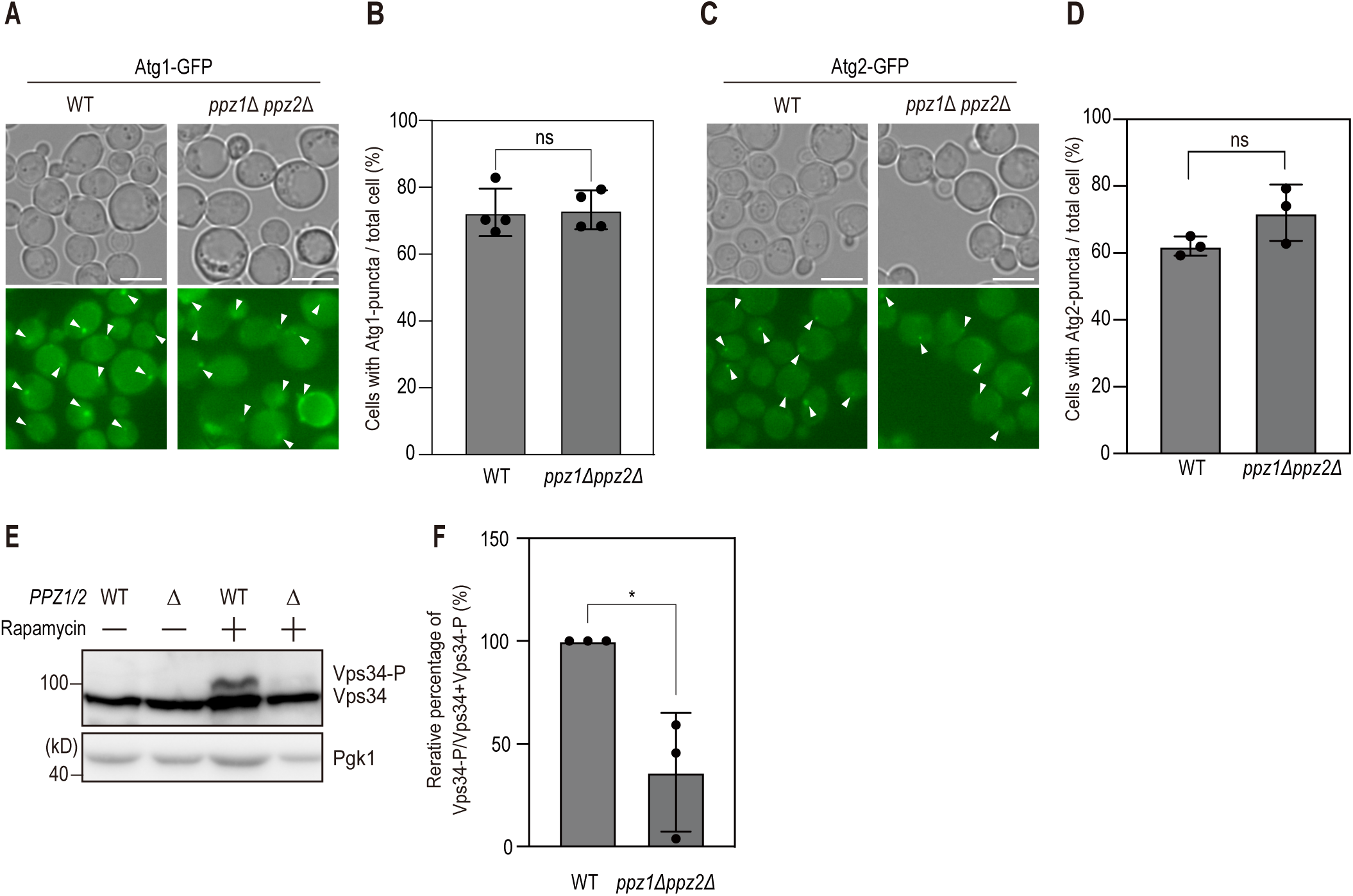
Deletion of *PPZ1* and *PPZ2* reduces Atg1 kinase activity. (A) Wild-type and *ppz1*Δ *ppz2*Δ cells expressing GFP-tagged Atg1 were examined by fluorescence microscopy after 3 h of rapamycin treatment. White arrowheads indicate puncta. Scale bars, 5 µm. (B) Quantification of the percentage of cells containing at least one Atg1–GFP punctum per cell. Data are presented as mean ± s.d. from independent cultures (n = 4). Unpaired t-test; *ns, P* ≥ 0.05. (C) Wild-type and *ppz1*Δ *ppz2*Δ cells expressing GFP-tagged Atg2 were examined by fluorescence microscopy after 3 h of rapamycin treatment. White arrowheads indicate puncta. Scale bars, 5 µm. (D) Quantification of the percentage of cells containing at least one Atg2-GFP punctum per cell. Data are presented as mean ± s.d. from independent cultures (n = 4). Unpaired t-test; *ns, P* ≥ 0.05. (E) Wild-type and *ppz1*Δ *ppz2*Δ cells were collected after 3 h of treatment with 0.2 µg/mL rapamycin. Lysates were analyzed by immunoblotting with anti-Vps34 and anti-Pgk1 antibodies. The graph shows mean ± s.d. from three independent experiments. Unpaired t-test: \**P* < 0.005 (F) The level of Vps34 phosphorylation was quantified and expressed as a ratio relative to total Vps34. Data are presented as mean values ± s.d. from three independent experiments (n = 3). Unpaired t-test: \**P* < 0.005

### Ppz1 and Ppz2 regulate autophagy through potassium transporters Trk1 and Trk2

While the induction of autophagy results in the recruitment of core Atg proteins to the PAS, we observed that Ppz1 localizes to multiple puncta at the periphery of the cell, even upon rapamycin treatment. (Supplemental Fig. 4A). This distinct spatial partitioning suggests that core Atg proteins, such as Atg1, are unlikely candidates for direct dephosphorylation by Ppz1/2. Ppz1 and Ppz2 are well-established negative regulators of the potassium transport activity mediated by Trk1 and Trk2 (hereafter Trk1/2), the primary potassium transporters in *S. cerevisiae* (Yenush et al., 2002). In agreement with a previous report (Yenush et al., 2005), we verified that GFP-tagged Trk1 localizes to the plasma membrane independently of rapamycin treatment (Supplemental Fig. 4B), placing it in the same subcellular compartment as Ppz1. To determine whether the aberrant potassium influx caused by the loss of Ppz1/2 underlies the observed autophagy defect, we generated a quadruple mutant by additionally deleting the major potassium transporters, *TRK1* and *TRK2*, in the *ppz1*Δ *ppz2*Δ background. Remarkably, the concomitant deletion of *TRK1/2* significantly suppressed the autophagy defect of *ppz1*Δ *ppz2*Δ cells (Fig. 4A). In line with this, GFP-Atg8 cleavage was also substantially restored in the quadruple-deletion strain, whereas GFP-Atg8 processing in the *trk1*Δ *trk2*Δ strain was unexpectedly elevated, even surpassing the levels observed in wild-type cells (Fig. 4B, 4C). The link between restricted potassium transport and enhanced autophagy is evident, but the precise molecular mechanism remains unclear. Furthermore, as TORC1 activity remained largely unaffected in the quadruple mutant, the rescue of autophagic activity in this genetic background is mediated by a mechanism acting downstream of TORC1 (Fig. 4D, Supplemental Fig. 4C).

**Figure 4.**
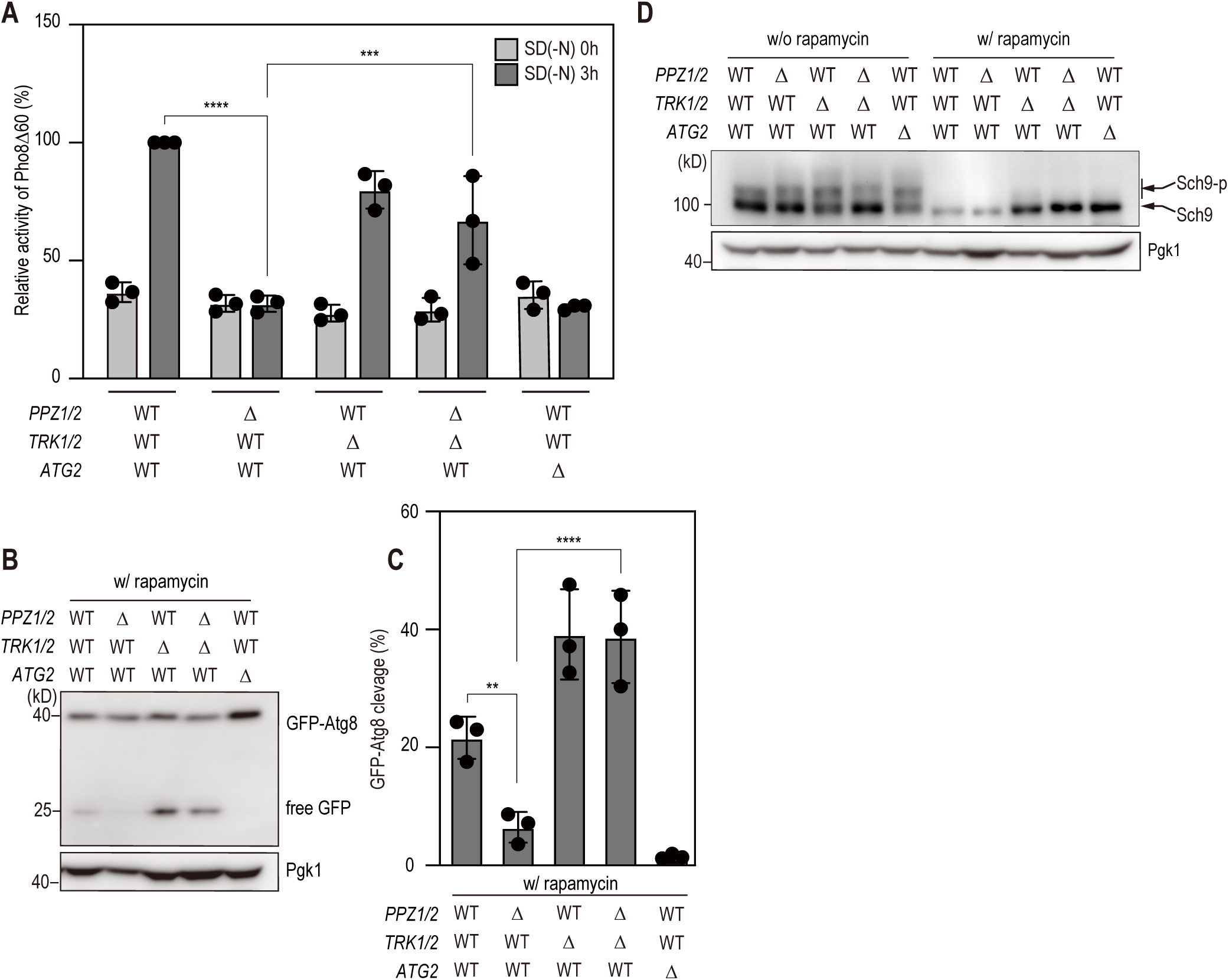
Potassium transport through Trk1 and Trk2 mediates Ppz1/2-dependent autophagy. (A) Wild-type cells and *ppz1*Δ *ppz2*Δ, *trk1*Δ *trk2*Δ, and *ppz1*Δ *ppz2*Δ *trk1*Δ *trk2*Δ mutants were incubated in YPD or SD-N medium for 3 h and subjected to an ALP assay at the indicated time points (n = 3). Tukey’s multiple comparison test: \*\*\*\**P* < 0.0001, \*\*\**P* < 0.001 (B) Cells were collected after 3 h of treatment with 0.2 µg/mL rapamycin. Lysates were analyzed by immunoblotting with anti-GFP and anti-Pgk1 antibodies. (C) Quantification of Fig. 4B. The ratio of free GFP to total GFP was quantified in each strain to assess autophagy flux. The graph shows mean values ± s.d. from three independent experiments. Tukey’s multiple comparison test: \*\*\*\**P* < 0.0001, \*\**P* < 0.01. (D) Cells were collected after 3 h of treatment with 0.2 µg/mL rapamycin. Lysates were analyzed by immunoblotting with anti-Sch9 and anti-Pgk1 antibodies.

### The suppression of potassium flux is indispensable for autophagy induction

Since Ppz1 and Ppz2 negatively regulate the potassium transporters Trk1 and Trk2 to restrict extracellular K^+^ influx, the loss of this inhibitory control in *ppz1*Δ *ppz2*Δ cells is predicted to lead to a concomitant rise in intracellular K^+^ concentrations. To investigate whether the elevated intracellular K^+^ concentration in *ppz1*Δ *ppz2*Δ cells directly underlies the autophagic defect, we performed GFP-Atg8 cleavage assays under potassium-limited conditions. For this purpose, we employed a modified SCD medium (designated SCD (–K^+^)) in which potassium phosphate was substituted with ammonium phosphate. This SCD (–K^+^) medium has a final potassium concentration of approximately 0.54 mM, which is less than 10% of this in standard SC media (7.87 mM). In line with previous reports, a low extracellular potassium concentration was sufficient to partially induce autophagy in wild-type cells without rapamycin treatment (Fig. 5A, B) (Rangarajan et al., 2020). Notably, this potassium limitation also triggered autophagy in *ppz1*Δ *ppz2*Δ cells, presumably because lowering the intracellular K^+^ pool alleviates the inhibitory constraint on the autophagic machinery. To determine whether low K^+^ could circumvent the *ppz1*Δ *ppz2*Δ autophagy defect under rapamycin-treated conditions, we cultured cells in potassium-limited medium; the GFP-Atg8 cleavage assay demonstrated a complete restoration of autophagic activity (Fig. 5A, B), indicating that the requirement for Ppz1/2 is bypassed when extracellular potassium is restricted. These findings collectively demonstrate that impaired autophagy in *ppz1*Δ *ppz2*Δ mutants is primarily attributable to the abnormal accumulation, implying that a reduction in these levels is essential for the induction of autophagy. To validate this model, we directly quantified intracellular K⁺ concentrations in the mutant strains. To determine K⁺ levels, we introduced a LAQUA twin K-11 ion meter (HORIBA) (Iwamoto et al., 2024). Notably, our measurements yielded values consistent with those reported in previous studies using inductively coupled plasma mass spectrometry (ICP-MS), confirming the reliability of this approach (Liu et al., 2019). In wild-type cells, intracellular K⁺ dropped by approximately 40% after 3 h of rapamycin treatment, from about 260 fg to 150 fg per cell, which is consistent with a prior report (Primo et al., 2017). In contrast, this decline was markedly attenuated in *ppz1*Δ *ppz2*Δ cells, with K⁺ levels decreasing by only approximately 20% (from about 290 fg to 220 fg per cell) following rapamycin treatment (Fig. 5C). In the *trk1*Δ *trk2*Δ strain, which effectively rescues the autophagic impairment of *ppz1*Δ *ppz2*Δ cells, intracellular K⁺ was maintained at constitutively low levels. Taken together, these findings provide compelling evidence for a model in which elevated intracellular K⁺ concentrations serve as a major inhibitory signal that blocks autophagy induction downstream of TORC1.

**Figure 5.**
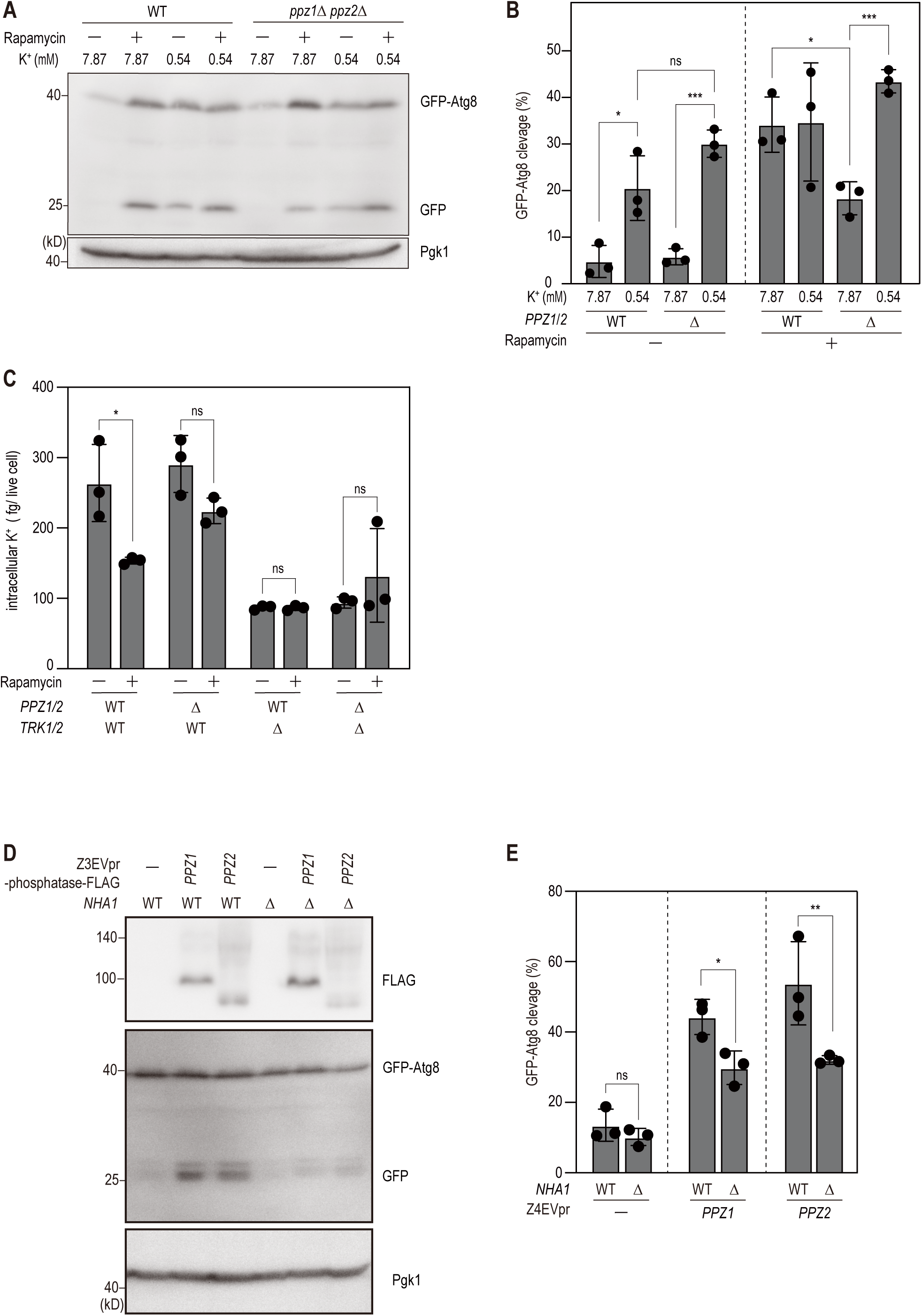
Potassium ion regulation is required for autophagy induction. (A) Cells were incubated in SC medium for 6 h and then transferred to one of the following conditions: SC with rapamycin, low-K^+^ medium, or low-K^+^ medium supplemented with rapamycin. Cells were further incubated for 3 h, and lysates were analyzed by immunoblotting with anti-GFP and anti-Pgk1 antibodies. SC medium contained 7.87 mM K^+^; low-K^+^ medium contained 0.54 mM K^+^. (B) Quantification of Fig. 5A. The graph shows mean values ± s.d. from three independent experiments. One-way ANOVA with Holm–Šidák correction: \*\*\**P* < 0.001, \**P* < 0.05*; ns, P* ≥ 0.05. (C) Cells were collected before (-) and after 3 h (+) of treatment with 0.2 µg/mL rapamycin. Intracellular potassium levels were quantified from the supernatant of lysates. Potassium amounts were normalized to cell viability and cell number. Data represent mean values ± s.d. from three independent experiments. Tukey’s multiple comparison test: \**P* < 0.05*; ns, P* ≥ 0.05. (D) Cells with indicated genotypes were collected after 3 h of treatment with β-estradiol. Lysates were analyzed by immunoblotting with anti-FLAG, anti-GFP, and anti-Pgk1 antibodies. (E) Quantification of Fig. 5D. The ratio of free GFP to total GFP was quantified in each strain to assess autophagy flux. The graph shows mean values ± s.d. from three independent experiments. Two-way ANOVA with Holm–Šidák correction for multiple comparisons to assess the effect of *NHA1* presence: \**P* < 0.01, \*\**P* < 0.05*; ns, P* ≥ 0.05.

Having demonstrated that aberrant K^+^ accumulation underlies the autophagic defect in the absence of Ppz1/2 phosphatases, we asked whether autophagy triggered by the overexpression of Ppz1 and Ppz2 is dependent on the modulation of potassium homeostasis. Nha1 is a plasma membrane Na^+^, K^+^/H^+^ antiporter that maintains cellular cation homeostasis by mediating the electrogenic efflux of K^+^ and Na^+^ in exchange for protons (Ariño et al., 2010). Given that intracellular K⁺ is expected to decrease dramatically upon overexpression of Ppz1 or Ppz2, we reasoned that deletion of *NHA1*—which limits K⁺ efflux—would attenuate this reduction in intracellular K^+^ and thereby abrogate the induction of autophagy. As predicted, the deletion of *NHA1* markedly diminished GFP-Atg8 processing in both Ppz1- and Ppz2-overexpressing cells (Fig. 5D, E). This epistatic interaction provides compelling evidence that the depletion of intracellular potassium represents a pivotal downstream event sufficient to trigger autophagy in response to Ppz1/2 hyperactivation.

## Discussion

In this study, we employed a gain-of-function screen to identify novel autophagy-related phosphatases that were not detected in previous loss-of-function screens by several groups (Harding et al., 1995; Thumm et al., 1994; Tsukada and Ohsumi, 1993), allowing us to identify the phosphatases Ppz1 and Ppz2 as regulators of autophagy induction. Compared to kinases, only approximately one third as many phosphatases have been identified in *S. cerevisiae* (127 kinases to 43 phosphatases), as well as humans and other organisms (Chen et al., 2017), suggesting that phosphatases must act on a broader range of substrates than their kinase counterparts. Ppz1 and Ppz2 were likely missed in earlier screens due to such functional redundancy (Cyert et al., 1991; Kim et al., 2011; Laviña et al., 2013) which causes partial, unclear phenotypes upon the deletion of single genes (Sakumoto et al., 2002); this effect is amplified by the paralogous nature of Ppz1 and Ppz2, as often observed in the yeast genome (Kellis et al., 2004). We were able to reproducibly detect an autophagy defect in the *ppz1*Δ *ppz2*Δ double deletion strain (Fig. 2A-C), highlighting the utility of gain-of-function screening to detect novel regulators of autophagy and other pathways. The Z3/4EV system (Louvion et al., 1993; McIsaac et al., 2013) allows rapid and dose-dependent induction of the gene of interest simply by adding β-estradiol. Moreover, for studies of nutrient dependent processes such as autophagy, this system is ideally suited over conventional induction methods that rely on the addition of copper or methionine, or on carbon source shifts—factors that themselves act as confounding modulators of autophagic activity.

Upon overexpression of Ppz1 or Ppz2, autophagy was induced even under nutrient rich conditions, in which autophagy is usually suppressed. Overexpression-based genetic screens rest on the premise that ectopic abundance of a signaling protein can constitutively activate a pathway independently of upstream stimuli. We observed that the phosphorylation states of the direct TORC1 substrates, Sch9 and Atg13, remained unchanged, indicating that the overexpression of Ppz1 or Ppz2 circumvented the necessity for TORC1 inactivation during autophagy induction (Fig. 1D, Supplemental Fig. 1A). Expression of an unphosphorylatable Atg13 mutant (Atg13-8SA) induces autophagy independently of TORC1 inactivation, similar to the effect observed with Ppz1 or Ppz2 overexpression (Codogno et al., 2012; Kamada et al., 2010). Although these results suggested that Atg13 might be a direct substrate for Ppz1 or Ppz2, our data argued against this possibility; Atg13 retained its hyperphosphorylated state even upon the overexpression of Ppz1 (Supplemental Fig. 1A). Furthermore, even in autophagy-defective *ppz1*Δ *ppz2*Δ cells, TORC1 activity was still reduced by rapamycin treatment (Fig. 2F). Together, these data suggest that Ppz1 or Ppz2 overexpression induces autophagy by dephosphorylating factors acting downstream of TORC1 and Atg13.

The majority of core Atg proteins are recruited to the pre-autophagosomal structure (PAS), which serves as the primary nucleation site for autophagosome biogenesis. The PAS is consistently positioned in proximity to the vacuole. By contrast, Ppz1 and Ppz2 reside at the plasma membrane under both nutrient-rich conditions and after treatment with rapamycin, despite the latter inducing autophagy. This spatial sequestration strongly suggests that the core Atg proteins are not a direct substrate of Ppz1/2; instead, these proteins likely operate through a distinct, peripheral mechanism, such as the modulation of intracellular ion homeostasis. Since Ppz1/2 are known regulators of the major K⁺ transporters Trk1 and Trk2 (Yenush et al., 2005), we investigated whether dysregulated K⁺ homeostasis provides a mechanistic link between these phosphatases and autophagy. Indeed, *ppz1*Δ *ppz2*Δ cells exhibited elevated intracellular K^+^ levels; however, autophagy was restored when K^+^ uptake was limited, either through the deletion of *TRK1* and *TRK2* or by culturing cells under low-K^+^ conditions. We also demonstrated that autophagy induction driven by Ppz1/2 overexpression is attenuated by the deletion of *NHA1*, a K^+^ efflux transporter. Overall, our data support the conclusion that a reduction in intracellular K^+^ levels serves as a potent trigger of autophagy, providing a molecular mechanism for the previously observed partial induction of autophagy under K^+^-limited conditions (Rangarajan et al., 2020). Integrating our findings from both Ppz1/2 overexpression and *ppz1*Δ *ppz2*Δ loss-of-function models, we propose a model wherein Ppz1/2-mediated suppression of Trk1/2 in response to nutrient starvation promotes a decline in intracellular K^+^ levels, thereby triggering autophagy.

In yeast, Trk1 and Trk2 mediate membrane-potential-dependent K^+^ uptake; this intracellular accumulation of K^+^ typically leads to membrane depolarization. This depolarization effect is buffered by the H^+^-ATPase Pma1, which facilitates ATP-dependent H^+^ efflux to stabilize the membrane potential (Madrid et al., 1998). Thus, K^+^ and H^+^ homeostasis are reciprocally linked, reflecting a stoichiometric trade-off between these two dominant monovalent cations within the cell. Within this framework, Ppz1/2-mediated inhibition of Trk1/2 likely shifts the balance toward reduced intracellular K^+^, whereas *ppz1*Δ *ppz2*Δ cells fail to recalibrate this shift and instead retain elevated intracellular K^+^. What are the molecular mechanisms underlying the autophagy defects arising from compromised K^+^ and H^+^ homeostasis? In our study, we observed that Atg1 kinase activity is markedly reduced in *ppz1*Δ *ppz2*Δ cells compared to wild-type cells (Fig. 3E, F). Although the direct functional links between Atg1 activity and K^+^/H^+^ concentrations that determine intracellular pH remain to be fully elucidated, the Atg1 complex exhibits high solubility *in vitro* at pH 7.0 but readily undergoes phase separation upon exposure to mildly acidic conditions (pH 6.0) (Noda et al., 2020). Thus, in *ppz1*Δ *ppz2*Δ cells, the synergistic effect of elevated K^+^ levels and a persistently neutral/alkaline cytosolic pH may collectively impede Atg1 phase separation, thereby suppressing its downstream kinase activity. Rapamycin treatment does not acidify cytosolic pH (Devare et al., 2020; Fujii et al., 2025), implying that a lack of H^+^ is unlikely to be the limiting factor. However, despite the elevated K^+^ and reduced H^+^ levels reported for *ppz1*Δ *ppz2*Δ cells (Yenush et al., 2005), PAS formation was still detectable in our study, suggesting that additional regulatory factors remain to be identified.

A particularly recent study in *Candida albicans* demonstrated that Ppz1 functions as a positive upstream regulator of TORC1 signaling under nutrient-rich conditions (Miao et al., 2025). In *ppz1*Δ/Δ mutants, a significant reduction in Rps6 phosphorylation is accompanied by the upregulation of autophagy-related genes, resulting in a characteristic “pseudo-starvation” state. Interestingly, the regulatory role of Ppz1 toward TORC1 appears to diverge between yeast species. Notably, the requirement for Ppz1 in TORC1 signaling diverges; it is obligatory for activation in *C. albicans* but dispensable in *S. cerevisia*e. The molecular basis for this interspecies divergence remains elusive, possibly reflecting evolutionary divergence to their distinct ecological niches. In this context, it would be highly intriguing to investigate whether *C. albicans ppz1*Δ/Δ mutants exhibit analogous autophagic defects during starvation. Such comparative studies would elucidate whether the role of Ppz1 in autophagy is a conserved ancestral function or is uniquely tethered to species-specific regulation of TORC1.

## Materials and methods

### Yeast strains and growth conditions

All yeast strains used in this study are listed in Table S2. All strains were derived from the W303-1A background (Thomas and Rothstein, 1989). Gene deletions and epitope tagging were performed using PCR-based methods as described previously (Baudin et al., 1993; Longtine et al., 1998). Successful tagging, integration, or deletion events were confirmed on selective plates and subsequently verified by PCR, fluorescence microscopy, or immunoblotting (Janke et al., 2004; Knop et al., 1999). All primers used in this study are listed in Table S3. Unless otherwise specified, genes of interest were expressed under the control of their endogenous promoters. Yeast strains were routinely cultured at 30°C in YPDA medium (1% yeast extract, 2% peptone, 2% dextrose, and 0.01% adenine sulfate). For nitrogen starvation experiments, cells were incubated in SD(-N) medium (0.17% yeast nitrogen base without amino acids and ammonium sulfate, supplemented with 2% glucose). For live-cell imaging, cells were cultured in synthetic complete (SC) medium (0.17% yeast nitrogen base without amino acids and ammonium sulfate, 0.5% ammonium sulfate, 2% glucose, 0.5% casamino acids, 0.002% tryptophan, 0.002% adenine, and 0.002% uracil). Autophagy was induced by treating cultures with 0.2 µg/mL rapamycin for the indicated times. For overexpression experiments, cells were treated with 2 µg/mL β-estradiol in YPDA or SC medium. For potassium limitation assays, yeast cultures were grown in low potassium synthetic medium (YNB without amino acids, ammonium sulfate, or potassium) supplemented with appropriate auxotrophic requirements. The pH of YNB-based media was adjusted to 5.8 with ammonium hydroxide before filtration.

### Plasmid construction

All plasmids used in this study are listed in Table S4. Plasmid amplification was performed in *Escherichia coli* DH5α cells under standard conditions. Site-directed mutations were introduced using PCR-based mutagenesis. Plasmid sequences were verified by Sanger sequencing (Genewiz, South Plainfield, NJ, USA).

### Fluorescence microscopy

Cells were cultured in synthetic complete dextrose (SC) medium and treated with 0.2 µg/mL rapamycin for 3 hours to induce autophagy. Cells were collected by centrifugation (600 × *g*, 2 min) and immediately subjected to fluorescence microscopy. Microscopy was performed using a Leica AF6500 fluorescence imaging system (Leica Microsystems, Wetzlar, Germany) mounted on a DMI6000B microscope equipped with an HCX PL APO 100×/1.40–0.70 oil-immersion objective lens and a xenon lamp (Leica Microsystems). Image acquisition and processing were carried out using LAS AF software (Leica Microsystems). Images were processed in Adobe Photoshop. The images were not manipulated other than contrast and brightness adjustments. Quantitative image analysis was subsequently performed using ImageJ/Fiji software, as described previously (Geng et al., 2008).

### Preparation of total yeast cell extract

Yeast cells (1-2 OD₆₀₀ units) were harvested and resuspended in 500 μl of cold ddH_2_O and 43 μl 100 % trichloroacetic acid, followed by brief vortexing and a 10-min incubation on ice. Precipitated proteins were pelleted by centrifugation (20,000 × *g*, 15 min, 4°C), and the supernatant was carefully removed. The resulting protein pellet was resuspended in HU buffer (200 mM sodium phosphate, pH 6.8, 8 M urea, 5% SDS, 0.1 mM EDTA, 0.005% bromophenol blue, and 15 mg/mL DTT) and transferred to a 1.5 mL screw-cap tube containing a small volume of 0.5-mm zirconia beads (Yasui Kikai, Osaka, Japan). Cells were disrupted using a FastPrep Cell Disruptor (MP Biomedicals, Irvine, CA, USA) and subsequently heated at 65°C for 15 min. The lysate was clarified by centrifugation (20,000 × *g*, 10 min, 25°C), and the supernatant was subjected to SDS–PAGE for protein separation.

### λ phosphatase treatment

Yeast cells (30–50 OD₆₀₀ units) were harvested by centrifugation (1,800 × g, 2 min, room temperature), resuspended in the remaining supernatant (0.5–1 mL), transferred to a 1.5 mL screw-cap tube, and collected by centrifugation (5,800 × g, 30 s, room temperature). The pellet was resuspended in 400 µL TAP-A buffer (50 mM Tris-HCl, pH 8.0, 150 mM NaCl, 10% glycerol, 1 mM DTT, 1 mM PMSF, 1 µM microcystin-LR, PhosSTOP, 1 mM EDTA, and Complete EDTA-free protease inhibitor cocktail) and transferred to a 1.5-mL screw-cap tube containing 100 µL of 0.5-mm zirconia beads (Yasui Kikai, Osaka, Japan). Cells were disrupted using a FastPrep Cell Disruptor (MP Biomedicals, Irvine, CA, USA) for three cycles of 30 s at speed 5.5 with 2 min on ice between cycles. Triton X-100 was added to a final concentration of 0.2%, followed by incubation for 10 min at 4°C. Lysates were clarified by centrifugation (17,800 × g, 10 min, 4°C), and the supernatant was collected. For immunoprecipitation, 5 µL anti-DYKDDDDK tag antibody magnetic beads (per sample) were washed three times with TAP-A buffer containing 0.2% Triton X-100 and incubated with clarified lysates for 1.5 h at 4°C with rotation to capture Ppz1-FLAG and ppz1D441N-FLAG. Beads were washed three times with TAP-A buffer containing 0.2% Triton X-100 and resuspended in 50 µL λ buffer (5 mM HEPES, pH 7.9, 2 mM MnCl₂, and 75 mM NaCl). λ phosphatase (5 µL) was added and incubated at 30°C for 20 min. Tubes were placed on a magnetic stand for 2 min and the supernatant was carefully removed. Bound proteins were eluted in HU buffer by heating at 65°C for 15 min, followed by SDS–PAGE and Western blotting.

### GFP-Atg8 cleavage assay

Cells were grown and subsequently treated with 0.2 µg/mL rapamycin for 3 h to induce autophagy. Equivalent cell amounts were collected for each condition and processed for immunoblotting as described in the previous section. Band intensities corresponding to free GFP and GFP–Atg8 were quantified using ImageJ/Fiji software. The rate of autophagic degradation was defined as the ratio of free-GFP to the sum of free-GFP and GFP-Atg8, expressed as a percentage (Araki et al., 2017). GFP cleavage (%) = free GFP / (free GFP + GFP–Atg8) × 100. Unless indicated otherwise, n = 3 independent biological replicates were performed; data are presented as mean ± s.d.

### ALP assay

Autophagic activity in yeast cells was quantified using an alkaline phosphatase (ALP) assay. The assay was performed following established protocols, and enzyme activity was measured spectrophotometrically (Araki et al., 2017).

### Intracellular K⁺ measurement

Yeast cultures (50 mL) were grown in synthetic complete (SC) medium (pH 5.8) to early-logarithmic phase (OD₆₀₀ ≈ 0.6). Cells were harvested by centrifugation, resuspended in 1 mL of distilled water, and incubated at 95°C for 10 min to release intracellular ions. The samples were centrifuged to remove cellular debris, and the potassium concentration in the resulting supernatant was quantified using a LAQUA twin K-11 ion meter (Horiba, Kyoto, Japan). Potassium standards (10–1000 ppm as K⁺) were prepared from a 1000-ppm K⁺ stock solution made by dissolving 1.908 g KCl per liter in ultrapure water. For routine calibration we used two points (100 and 500 ppm) according to the manufacturer’s instructions. For linearity assessment, solutions at 50, 100, 150, 200, 250, 300, 350, 400, 450, 500, 750, and 1000 ppm were measured in triplicate at room temperature (25°C). The sensor was rinsed with ultrapure water and blotted between measurements. Readings were recorded after stabilization. Data were analyzed by ordinary least squares on the mean of triplicates, and residuals and regression coefficients (slope, intercept, and R²) are reported in the corresponding figures. Intracellular potassium content per cell was subsequently determined by normalizing the measured potassium concentration to cell number. Cell numbers were estimated from OD₆₀₀ values using a predetermined conversion factor (Supplemental Fig. 5A, B) and verified by direct counting with a hemocytometer. To assess viability, cells were stained with phloxine B, and live/dead ratios were determined by fluorescence microscopy. Potassium content was normalized to the number of viable cells to obtain intracellular potassium content per living cell.

### Statistical analyses

Statistical analyses were performed using an unpaired, two-tailed Student’s *t* test. For multiple comparisons, one-way ANOVA followed by Tukey’s post hoc multiple-comparisons test was used. Differences were considered statistically significant at *P* < 0.05 (*), *P* < 0.01 (**), *P* < 0.001 (***), and *P* < 0.0001 (****). “ns” denotes not significant.

## Declaration of Generative AI in the writing process

ChatGPT (OpenAI, San Francisco, CA, USA) was used to assist in the literature survey and for initial English editing. All resulting text was subsequently checked and revised. The authors take full responsibility for the final content of the paper.

## Acknowledgements

We are grateful to Dr. Yoshinori Ohsumi for the kind gifts of antibodies and plasmids. We thank A. May for comments on the manuscript. This work was supported by Grants-in-Aid for Scientific Research from the Ministry of Education, Culture, Sports, Science and Technology (MEXT) of Japan to T.N (22H04647, 23H02475) and Y.A(24K09455), and Ohsumi Frontier Science Foundation to T.N.. N.M. was supported by JST SPRING (JPMJSP2138).

## Author contributions

Conceptualization: Y.A. and N.M.; Formal analysis: N.M.; Investigation: N.M. and Y.A.; Resource: S.A. and S.A.; Writing – original draft: N.M. and Y.A.; Writing – review and editing: N.M. and Y.A., with input from all authors; Visualization: N.M.; Supervision: T.N.; Funding acquisition: N.M., Y.A., and T.N.

## Data and resource availability

All relevant data and details of resources can be found within the article and its supplementary information.

---

**Supplemental Table S1. Results of overexpressing each of the 39 protein phosphatases encoded by the *Saccharomyces cerevisiae* genome**

The notation “+” indicates GFP-Atg8 cleavage or an increase in GFP-Atg8, whereas “–” denotes no cleavage or no increase. Since elevated total GFP-Atg8 levels can result from TORC1 suppression upon overexpression, strains exhibiting stable GFP-Atg8 levels were selected for further analysis.

**Supplemental Figure S1. Overexpression of Ppz1 does not reduce Atg13 phosphorylation. ppz1D441N is phosphorylated**

(A) Atg13-GFP-tagged cells with or without Ppz1-FLAG were collected after 3 h of treatment with β-estradiol, rapamycin, or both. Lysates were analyzed by immunoblotting with anti-FLAG, anti-GFP, and anti-Pgk1 antibodies.

(B) *ppz1D441N* cells were treated with 2 µg /mL β-estradiol for 3 h. Lysates were treated with λ phosphatase and analyzed by immunoblotting with anti-FLAG antibodies.

**Supplemental Figure S2. Loss of *PPZ1* and *PPZ2* impairs autophagy dependent accumulation of Atg1 and Atg8 in the vacuoles**

(A) GFP-tagged-Atg1 cells with or without *PPZ1* and *PPZ2* were examined by fluorescence microscopy after 3 h of rapamycin treatment. Scale bars, 5 µm.

(B) GFP-tagged-Atg8 cells with or without *PPZ1* and *PPZ2* were examined by fluorescence microscopy after 3 h of rapamycin treatment. Scale bars, 5 µm.

**Supplemental Figure S3. Loss of *PPZ1* and *PPZ2* does not affect Atg2 PAS recruitment**

(A) GFP-tagged-Atg2 cells with or without *PPZ1* and *PPZ2* were examined by fluorescence microscopy after 3 h of rapamycin treatment.

(B) Atg2-GFP puncta colocalizing with Atg17-mCherry were quantified; the cytosolic GFP signal of the same cell was used as background. Unpaired t-test; *ns, P ≥* 0.05.

**Supplemental Figure S4. The localization of Ppz1 and Trk2. Rapamycin induces Atg13 dephosphorylation in *ppz1*Δ *ppz2*Δ, *trk1*Δ *trk2*Δ, and *ppz1*Δ *ppz2*Δ *trk1*Δ *trk2*Δ mutants to levels comparable to wild type**

(A) GFP-tagged-Ppz1 cells were examined by fluorescence microscopy before and after 3 h of rapamycin treatment.

(B) GFP-tagged-Trk1 cells were examined by fluorescence microscopy before and after 3 h of rapamycin treatment.

(C) Wild-type cells and *ppz1*Δ *ppz2*Δ, *trk1*Δ *trk2*Δ, and *ppz1*Δ *ppz2*Δ *trk1*Δ *trk2*Δ mutants were collected before and after 3 h of treatment with 0.2 µg/mL rapamycin. Lysates were analyzed by immunoblotting with anti-Atg13 and anti-Pgk1 antibodies.

**Supplemental Figure S5. Calibration curve and residual plot for potassium detection (50–500 ppm)**

(A) Calibration curve for potassium using the HORIBA LAQUA twin K-11 sensor across 50–500 ppm (n = 3 per level). Each point represents the mean of triplicate measurements. Linear regression yielded y = −2.578 + 1.01895x with R² = 0.99932.

(B) Pointwise residuals (Measured − Predicted) versus target concentration with a horizontal reference line at 0. Residuals showed no curvature or heteroscedastic pattern. RMSE = 3.24 ppm; maximum relative error = 3.33% (at 100 ppm).

**Table S1.** Screening result of phosphatases.

| phosphatase | GFP-Atg8<br>cleavage | GFP-Atg8<br>increase | phosphatase | GFP-Atg8<br>cleavage | GFP-Atg8<br>increase |
| --- | --- | --- | --- | --- | --- |
| *PPZ1* | + | - | *PPH3* | - | - |
| *PPZ2* | + | - | *SIT4* | - | - |
| *SSU72* | - | - | *YVH1* | - | - |
| *SDP1* | - | - | *PPH21* | - | - |
| *OCA1* | - | - | *PPG1* | - | - |
| *PTC1* | - | - | *PPH22* | - | - |
| *GLC7* | - | - | *CDC14* | - | - |
| *PTC7* | - | - | *PSR2* | - | - |
| *PTP1* | - | - | *PSR1* | - | - |
| *PTC6* | - | - | *TEP1* | - | - |
| *PTC2* | - | - | *PPT1* | - | - |
| *NEM1* | - | - | *PTC4* | + | + |
| *PTC3* | - | - | *CNA1* | - | - |
| *MSG5* | - | - | *MIH1* | - | - |
| *PPQ1* | - | - | *PTC5* | - | - |
| *PTP2* | - | - | *CMP2* | - | - |
| *YCH1* | + | + | *FCP1* | - | - |
| *LTP1* | - | - | *PPS1* | - | - |
| *OCA2* | - | - | *PTP3* | - | - |
| *SIW14* | - | - |  |  |  |

**Table S2.**
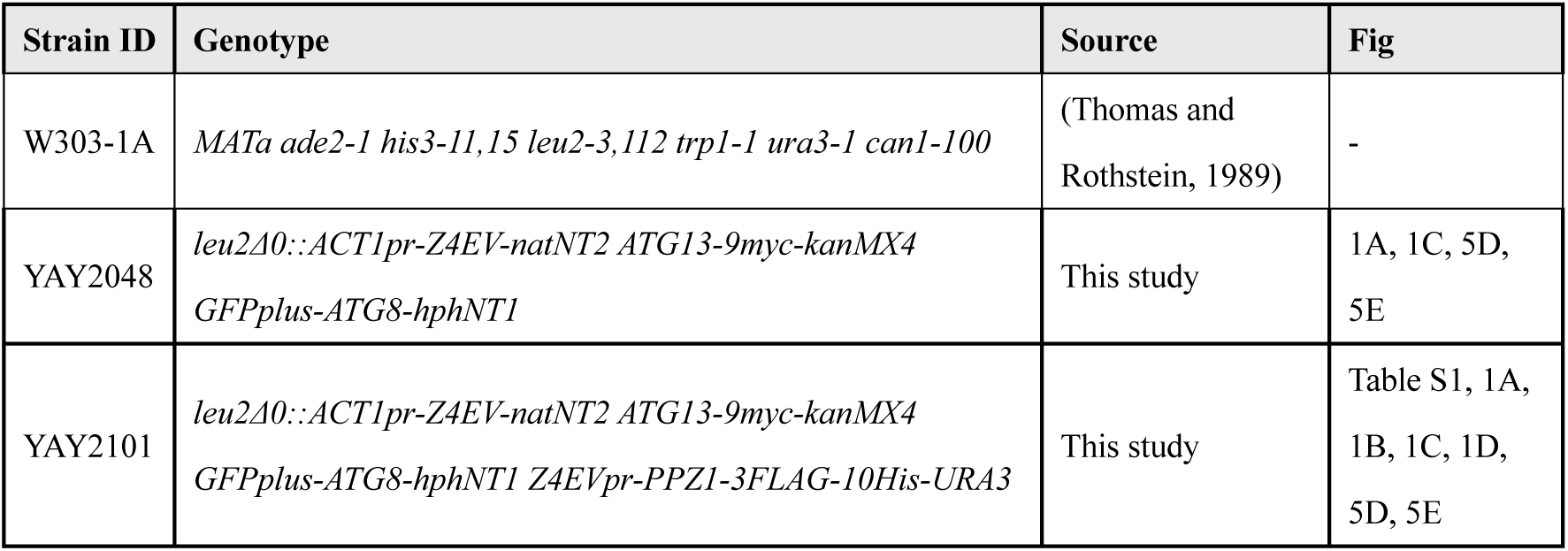
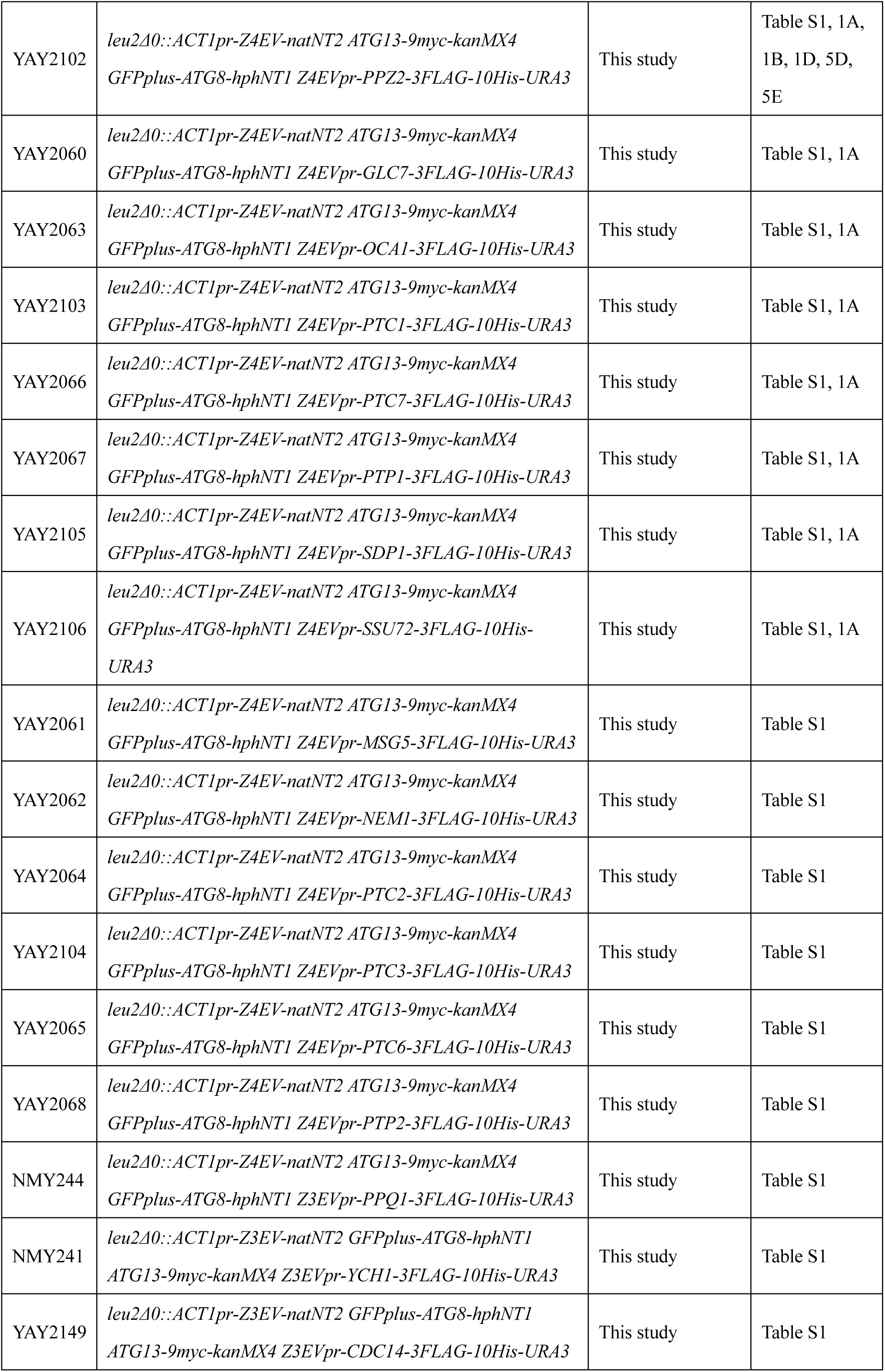
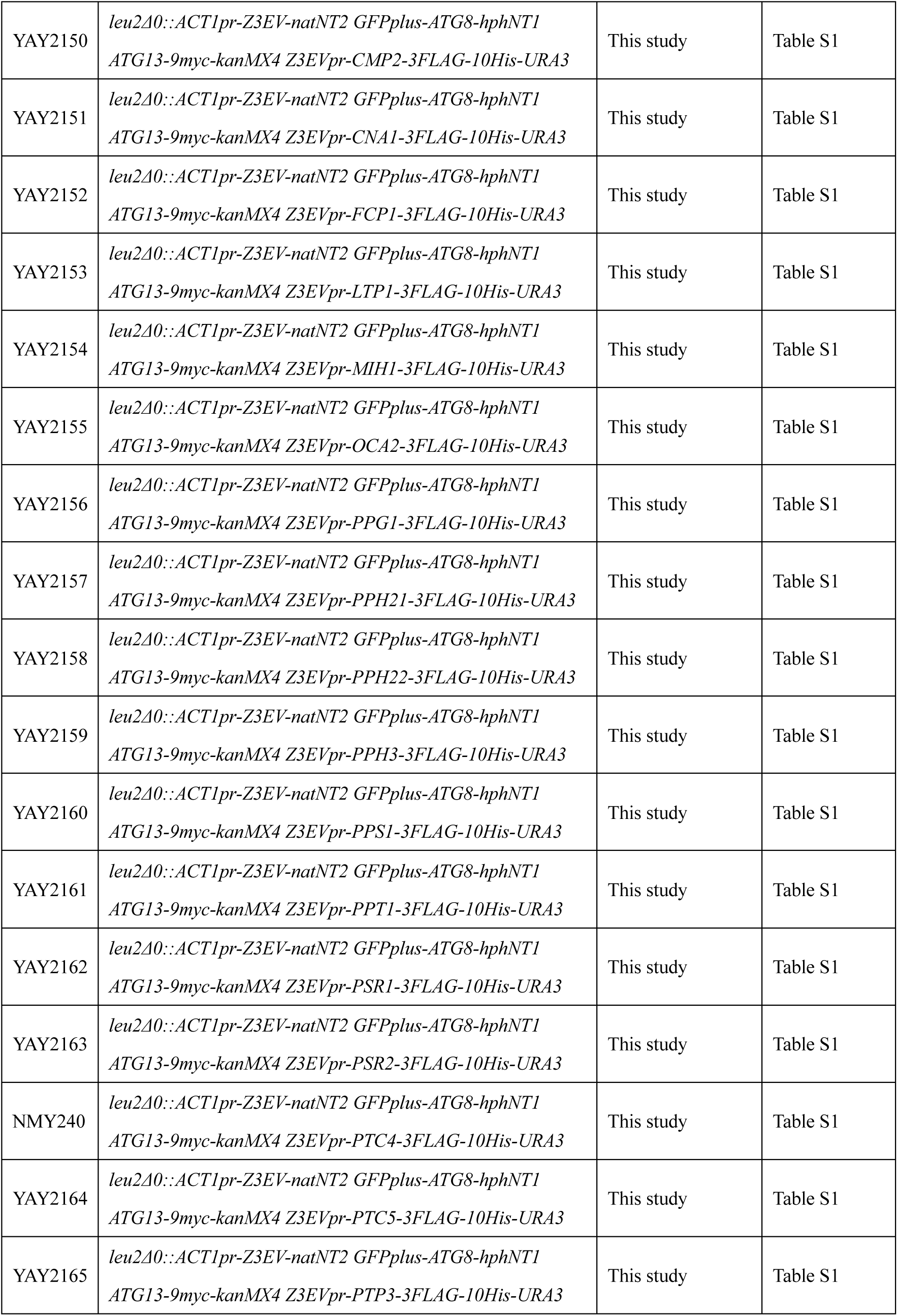
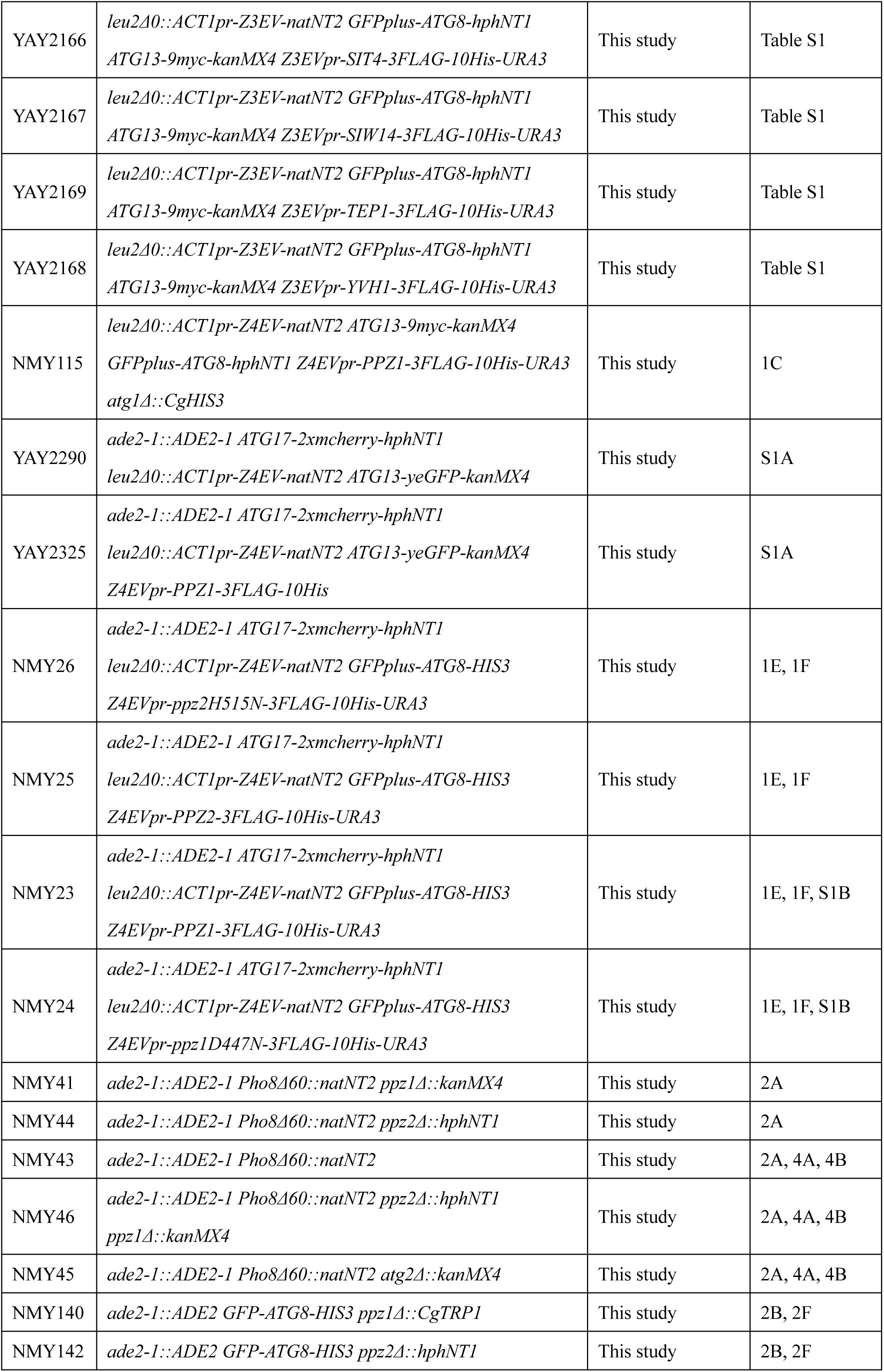
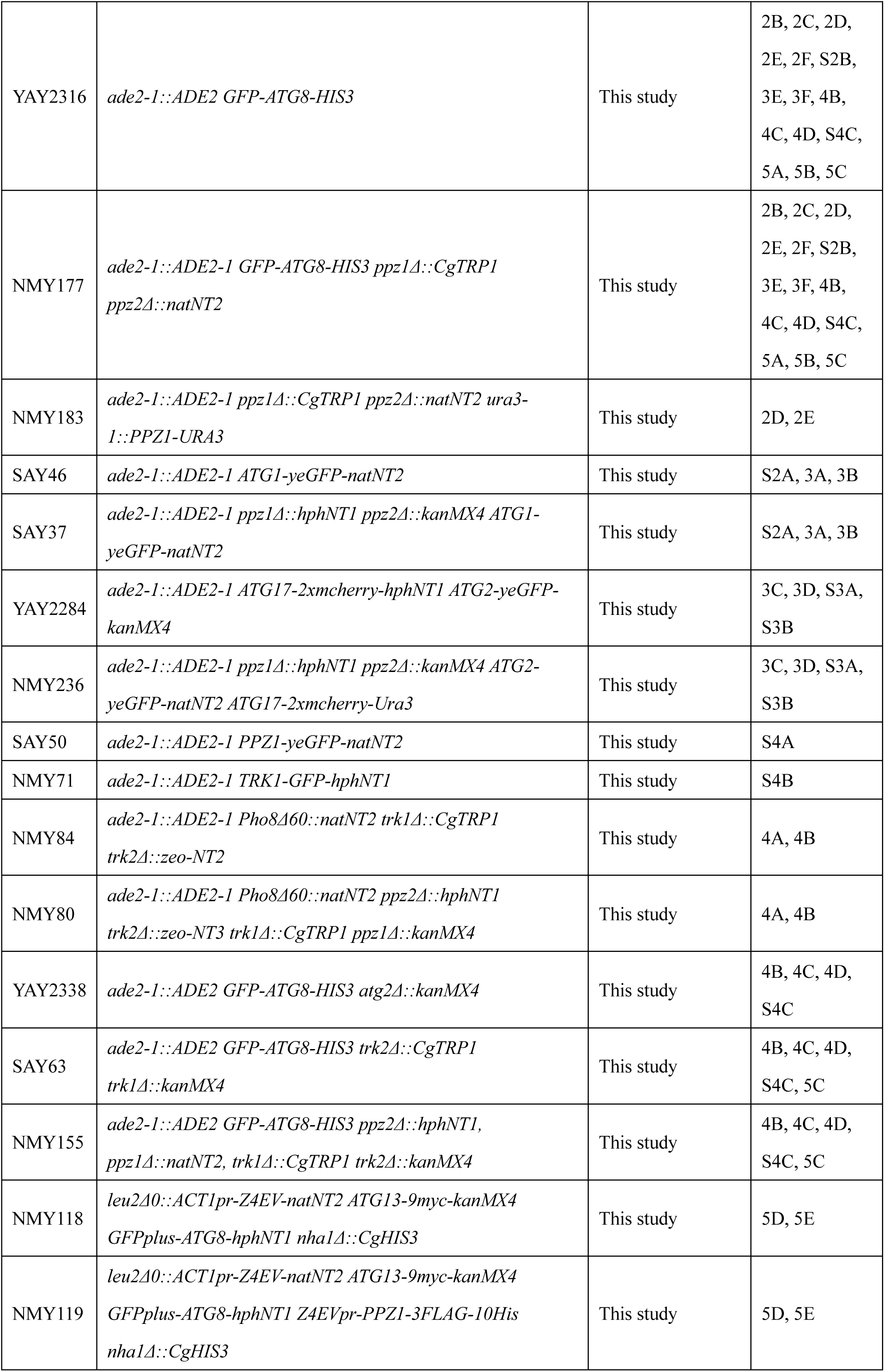
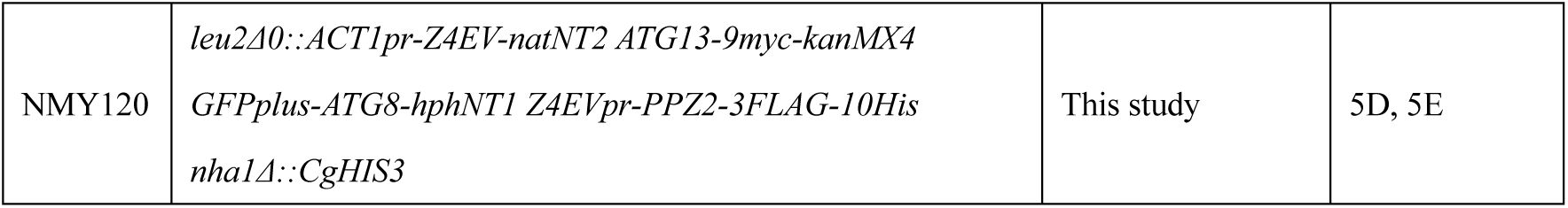
Strains used in this study.

**Table S3.**
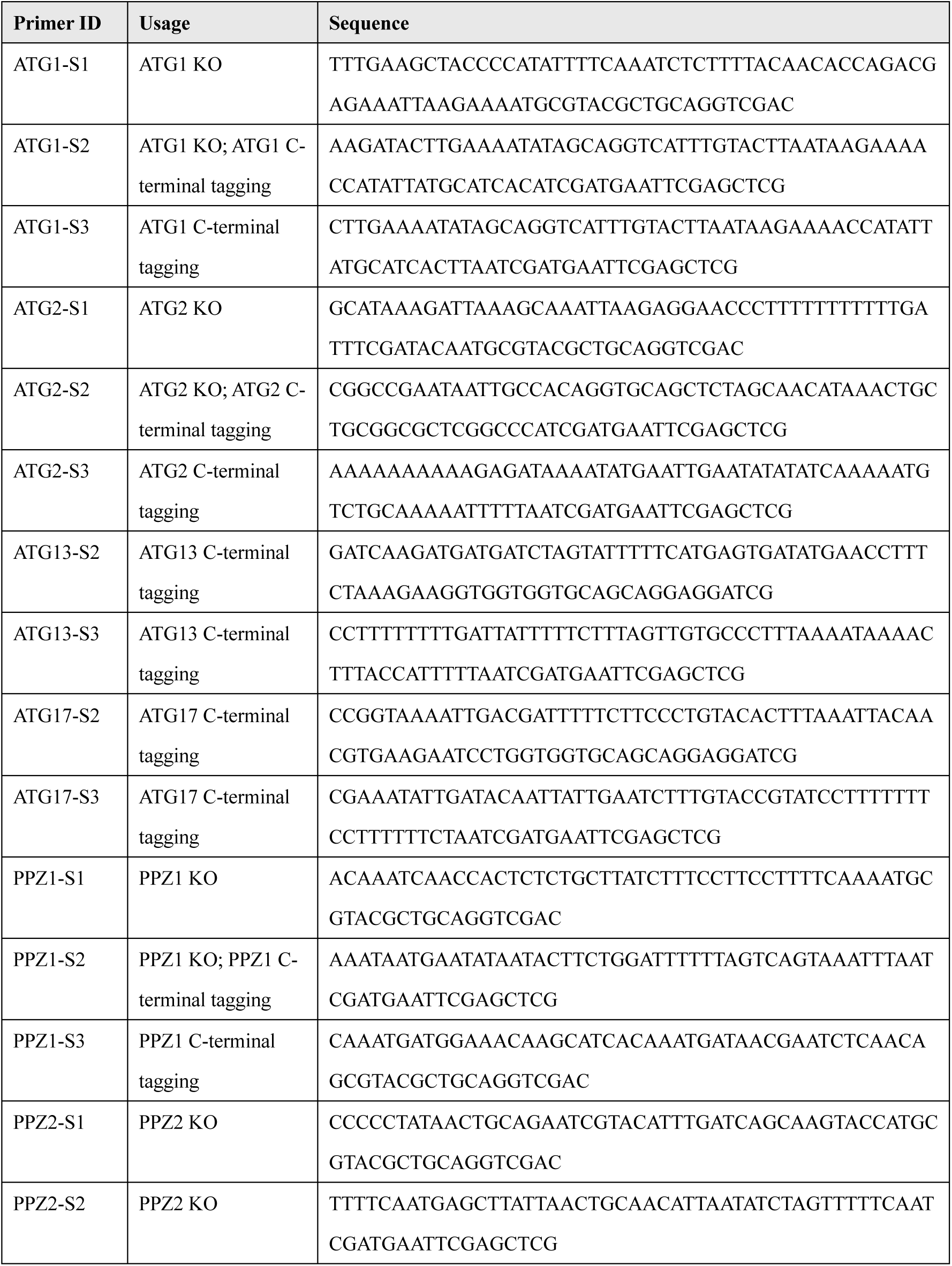
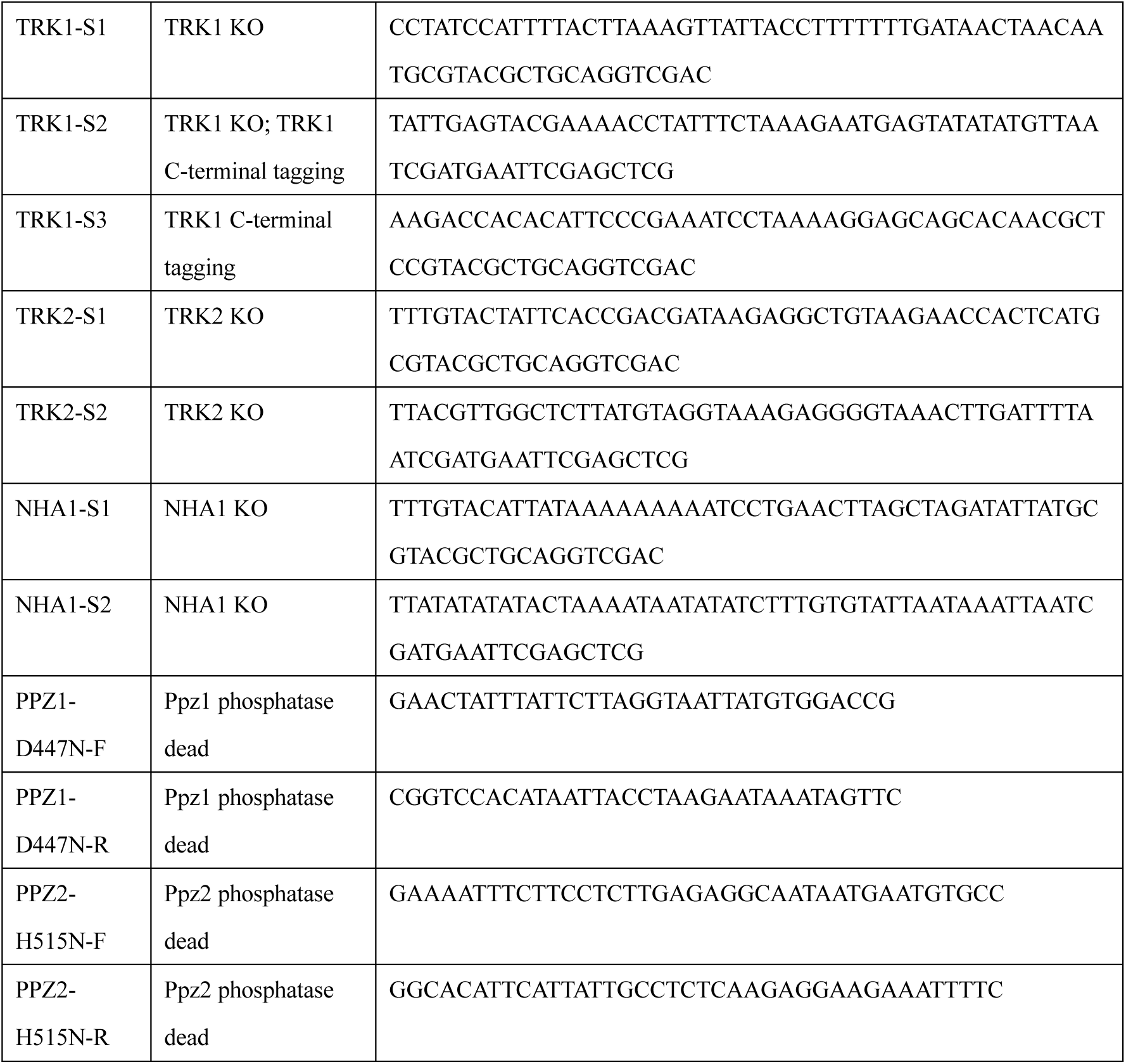
Oligo primers used in this study.

**Table S4.**
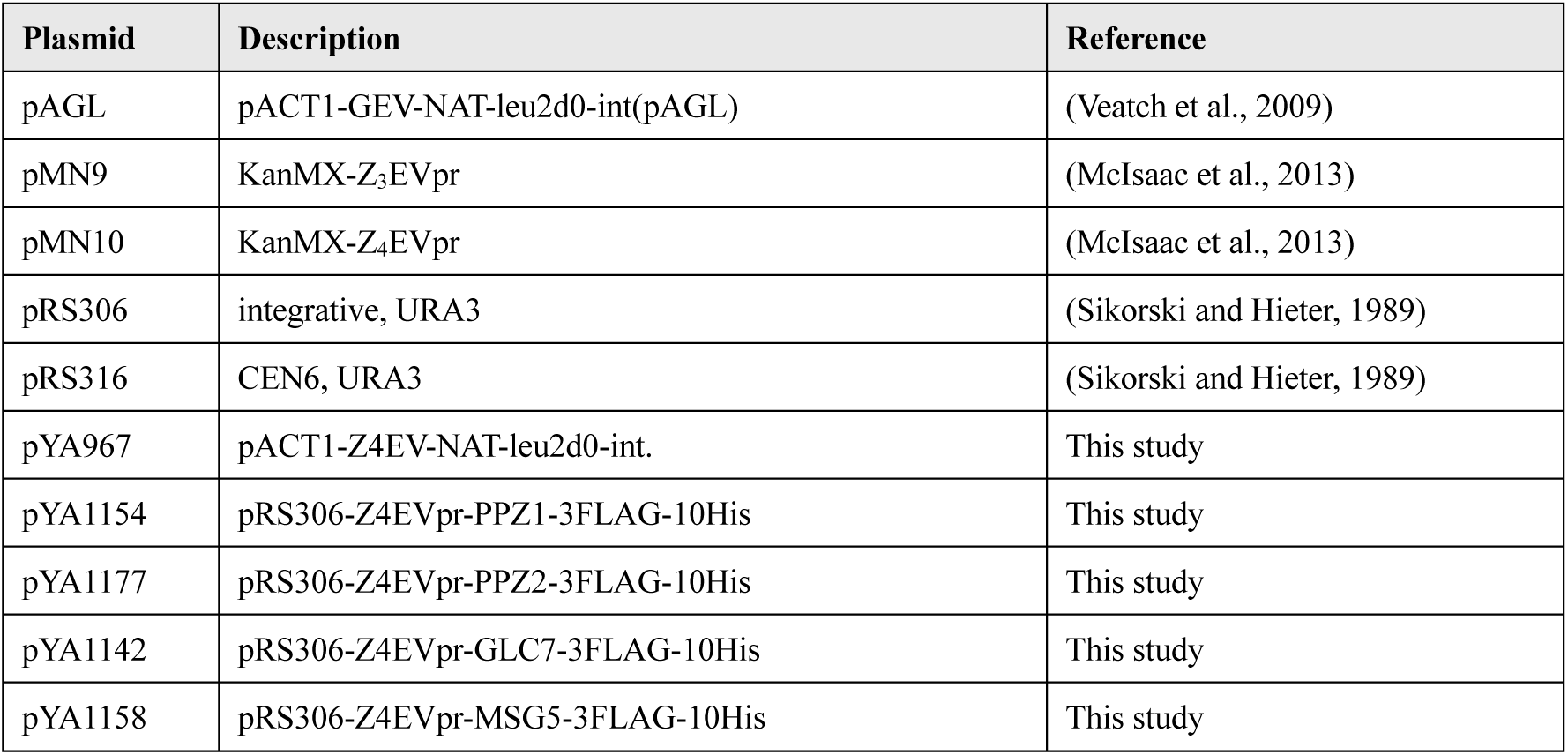
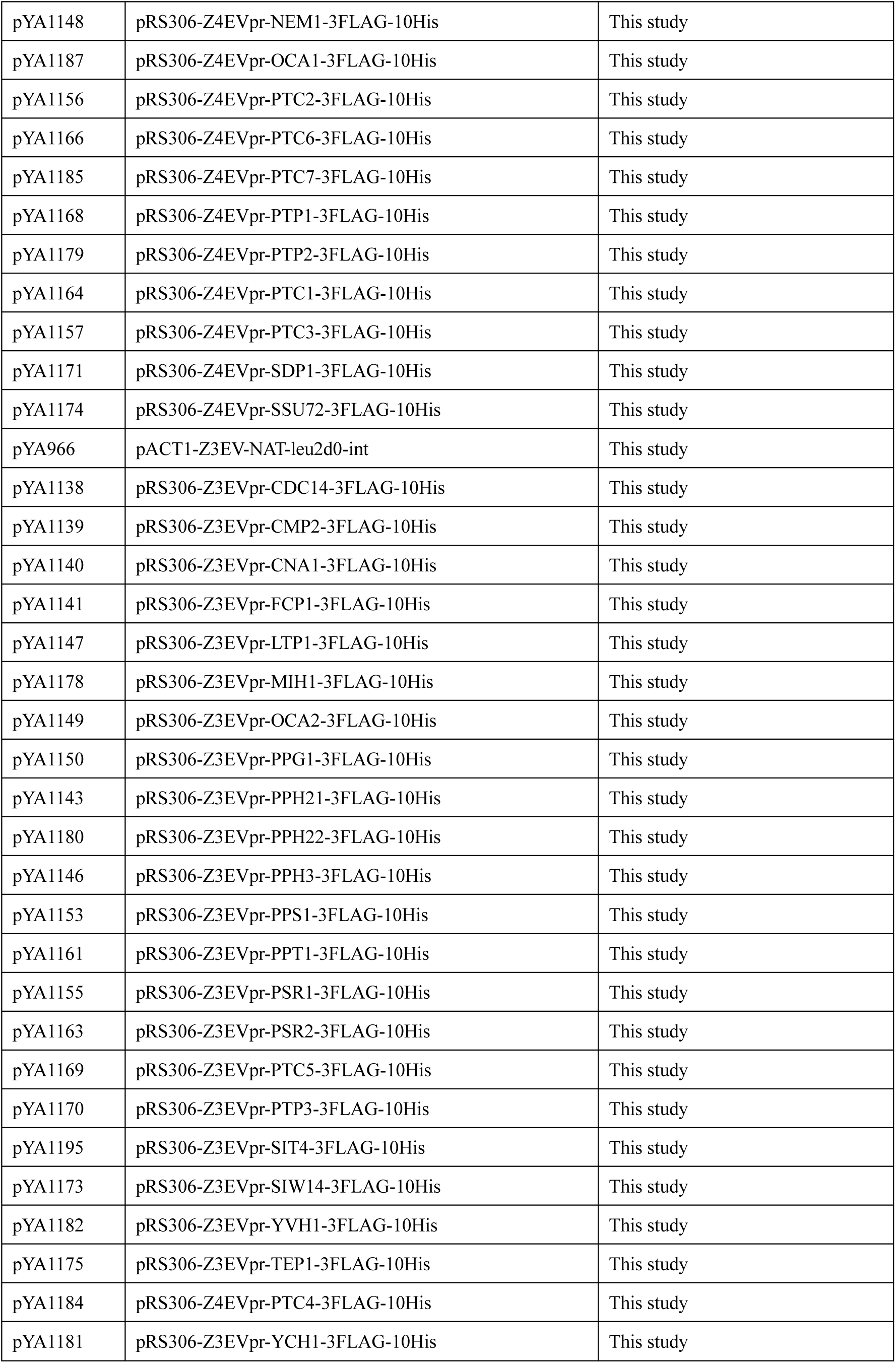
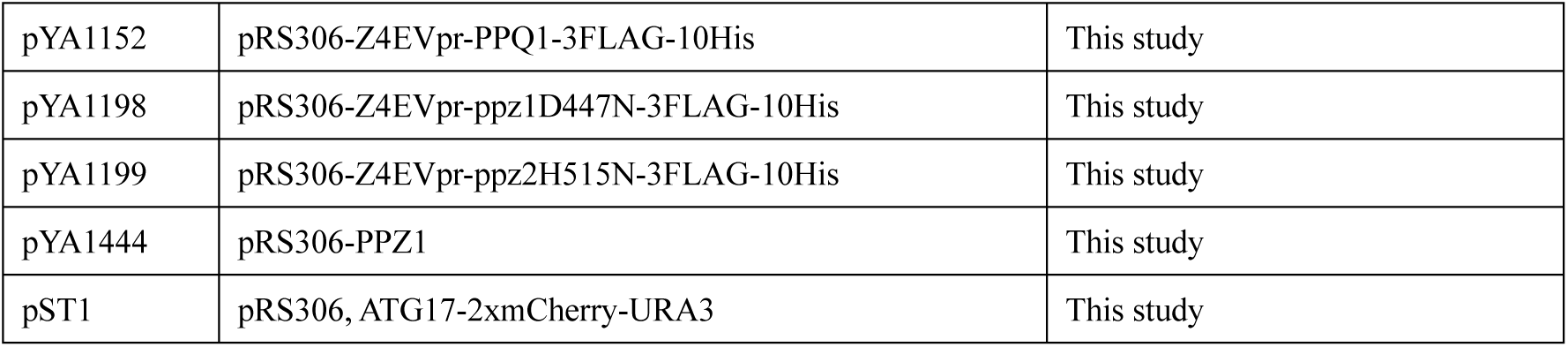
Plasmids used in this study.

**Table S5.** Reagents used in this study.

| REAGENT or RESOURCE | SOURCE | IDENTIFIER Catalog No.;RRID; Lot# | DILUTIONS |
| --- | --- | --- | --- |
| **Antibodies** |  |  |  |
| anti-phospho-Sch9-T737 antibody | Lab stock | N/A | 1/1000 |
| anti-Sch9 antibody | Lab stock | N/A | 1/1000 |
| HRP-conjugated goat anti-mouse IgG antibody | Jackson ImmunoResearch Laboratories, Inc. (West Grove, PA, USA) | 115-035-003; RRID: AB_10015289; Lot#176400 | 1/5000 |
| HRP-conjugated goat anti-rabbit IgG antibody | Jackson ImmunoResearch Laboratories, Inc. (West Grove, PA, USA) | 111-035-003; RRID: AB_2313567; Lot#176400 | 1/5000 |
| Rabbit anti-API antibody | Lab stock (originally gifted from Dr. Ohsumi Lab) | N/A | 1/2000 |
| Mouse monoclonal anti-FLAG M2 antibody | Sigma-Aldrich (St Louis, MO, USA) | F1804; RRID: AB_262044; Lot#SLBG5673V | 1/2000 |
| Mouse monoclonal anti-GFP antibody | Roche (Basel, Switzerland) | 11814460001; RRID: AB_390913; Lot#14442000 | 1/2000 |
| Mouse monoclonal anti-PGK1 antibody | Thermo Fisher Scientific (Waltham, MA, USA) | 459250; RRID: AB_2532235; Lot#WG3323356 | 1/10000 |
| Rabbit anti-Atg13 antibody | Lab stock (originally gifted from Dr. Ohsumi Lab) | N/A | 1/1000 |
| Rabbit anti-Vps34 antibody | Lab stock (originally gifted from Dr. Ohsumi Lab) | N/A | 1/1000 |

| REAGENT or RESOURCE | SOURCE | IDENTIFIER |
| --- | --- | --- |
| **E. coli** |  |  |
| *E. coli* DH5α competent cells | BioElegen Technology Co., Ltd.<br>(Taichung, Taiwan) | DH01-20 |

| REAGENT or RESOURCE | SOURCE | IDENTIFIER<br>(catalog No.) |
| --- | --- | --- |
| **Chemicals** |  |  |
| 1-Naphthyl phosphate | Sigma-Aldrich (St Louis, MO, USA) | N7255-5 |
| 30 (w/v) % Acrylamide/Bis Mixed Solution | Nacalai Tesque (Kyoto, Japan) | 06144-05 |
| 6-Aminohexanoic acid | FUJIFILM Wako (Osaka, Japan) | 012-09645 |
| Acetic acid | FUJIFILM Wako (Osaka, Japan) | 017-00256 |
| Adenine sulfate | FUJIFILM Wako (Osaka, Japan) | 010-19612 |
| Agarose | Nacalai Tesque (Kyoto, Japan) | 01208-85 |
| Albumin standard (BSA) | Thermo Fisher Scientific<br>(Waltham, MA, USA) | 23209 |
| Ammonium hydroxide | FUJIFILM Wako (Osaka, Japan) | 013-23355 |
| Ammonium persulfate (APS) | FUJIFILM Wako (Osaka, Japan) | 016-20501 |
| Ammonium sulfate | Nacalai Tesque (Kyoto, Japan) | 02619-15 |
| Anti DYKDDDDK tag antibody magnetic beads | FUJIFILM Wako (Osaka, Japan) | 017-25151 |
| Bacto peptone | BD Biosciences (San Jose, CA, USA) | 211677 |
| Bromophenol blue | Sigma-Aldrich (St Louis, MO, USA) | B0126 |
| Casamino acids | BD Biosciences (San Jose, CA, USA) | 223050 |
| Chemi-Lumi One Ultra | Nacalai Tesque (Kyoto, Japan) | 11644-40 |
| Complete EDTA-free protease inhibitor cocktail | Sigma-Aldrich (St Louis, MO, USA) | 11873580001 |
| D(+)-Glucose | FUJIFILM Wako (Osaka, Japan) | 043-31163 |
| Disodium dihydrogen ethylenediaminetetraacetate dihydrate | Nacalai Tesque (Kyoto, Japan) | 15130-95 |
| Disodium hydrogenphosphate dodecahydrate | FUJIFILM Wako (Osaka, Japan) | 196-02835 |
| DMSO | Sigma-Aldrich (St Louis, MO, USA) | D8418 |
| DTT | FUJIFILM Wako (Osaka, Japan) | 042-29222 |
| EtBr solution | NIPPON GENE (Tokyo, Japan) | 315-90051 |
| Ethanol | Nacalai Tesque (Kyoto, Japan) | 14713-95 |
| Glycine | FUJIFILM Wako (Osaka, Japan) | 073-00737 |
| KCl | Sigma-Aldrich (St Louis, MO, USA) | P9541 |
| KOD One PCR Master mix | Toyobo Co., Ltd., Osaka, Japan | KMM-101 |
| L-Tryptophan | Sigma-Aldrich (St Louis, MO, USA) | 93659 |
| Lithium acetate | FUJIFILM Wako (Osaka, Japan) | 127-01545 |
| Magnesium chloride hexahydrate | Nacalai Tesque (Kyoto, Japan) | 20908-65 |
| Methanol | FUJIFILM Wako (Osaka, Japan) | 139-01827 |
| Microcystin-LR | FUJIFILM Wako (Osaka, Japan) | 136-12241 |
| Morinaga skim milk | Morinaga milk (Tokyo, Japan) | N/A (commercial product) |
| Phloxine B | Sigma-Aldrich (St Louis, MO, USA) | P2759 |
| PhosSTOP | Roche (Basel, Switzerland) | 4906837001 |
| PMSF | FUJIFILM Wako (Osaka, Japan) | 164-12181 |
| Polyethylene glycol 3350 (PEG 3350) | FUJIFILM Wako (Osaka, Japan) | 1546547 |
| Protein Assay BCA kit | Nacalai Tesque (Kyoto, Japan) | 06385-00 |
| Rapamycin | LKT Laboratories, Inc. (St Paul, MN, USA) | R-5000 |
| Sodium chloride | FUJIFILM Wako (Osaka, Japan) | 195-01663 |
| Sodium hydroxide | FUJIFILM Wako (Osaka, Japan) | 194-18865 |
| Sodium lauryl sulfate | Nacalai Tesque (Kyoto, Japan) | 08933-05 |
| Sorbitol (D-glucitol) | Nacalai Tesque (Kyoto, Japan) | 32021-24 |
| TEMED (*N,N,N',N'*-tetramethyl-ethylene diamine) | FUJIFILM Wako (Osaka, Japan) | 205-66313 |
| Translucent K<sup>+</sup>-free yeast nitrogen base without amino acids, ammonium sulfate and potassium | Formedium Ltd. (Norfolk, UK) | CYN75 |
| Trichloroacetic acid | FUJIFILM Wako (Osaka, Japan) | 200-08085 |
| Tris (hydroxymethyl) aminomethane | Nacalai Tesque (Kyoto, Japan) | 35406-91 |
| Triton X-100 | FUJIFILM Wako (Osaka, Japan) | 581-81705 |
| Tween 20 | FUJIFILM Wako (Osaka, Japan) | 166-21115 |
| Uracil | FUJIFILM Wako (Osaka, Japan) | 212-00062 |
| Urea | FUJIFILM Wako (Osaka, Japan) | 219-00175 |
| Yeast extract | BD Biosciences (San Jose, CA, USA) | 212750 |
| Yeast nitrogen base without amino acids and ammonium sulfate | BD Biosciences (San Jose, CA, USA) | 233520 |
| Zinc sulfate heptahydrate | FUJIFILM Wako (Osaka, Japan) | 268-00405 |
| β-estradiol | FUJIFILM Wako (Osaka, Japan) | 052-04041 |
| $\lambda$ protein phosphatase | New England Biolabs (Ipswich, MA, USA) | P0753S |

| REAGENT or RESOURCE | SOURCE |
| --- | --- |
| **Software and algorithms** |  |
| GraphPad Prism | GraphPad Software, LLC (San Diego, CA, USA) |
| Illustrator (Adobe Inc., San Jose, CA, USA) | Adobe Inc. (San Jose, CA, USA) |
| ImageJ | National Institutes of Health (Bethesda, MD, USA) |
| Photoshop | Adobe Inc. (San Jose, CA, USA) |

