## Supplemental Figure S1-S5 for "Autophagy induction requires the suppression of potassium influx mediated by phosphatases"

### Supplymental Fig1(Supp of Fig. 1)

**A**

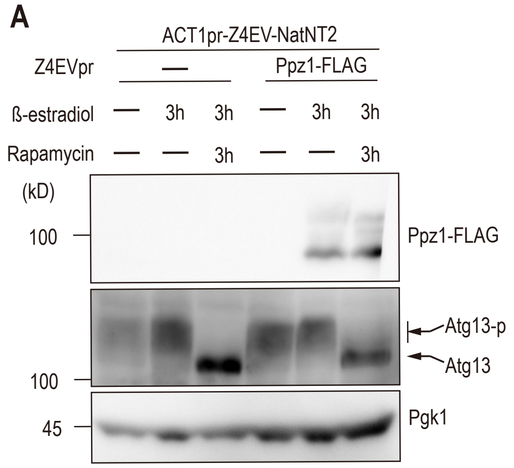


**B**

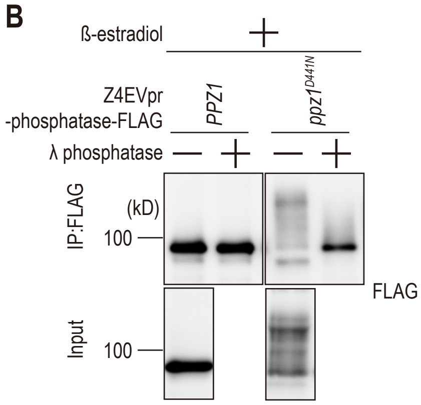

Supplymental Fig2(Supp of Fig. 2)

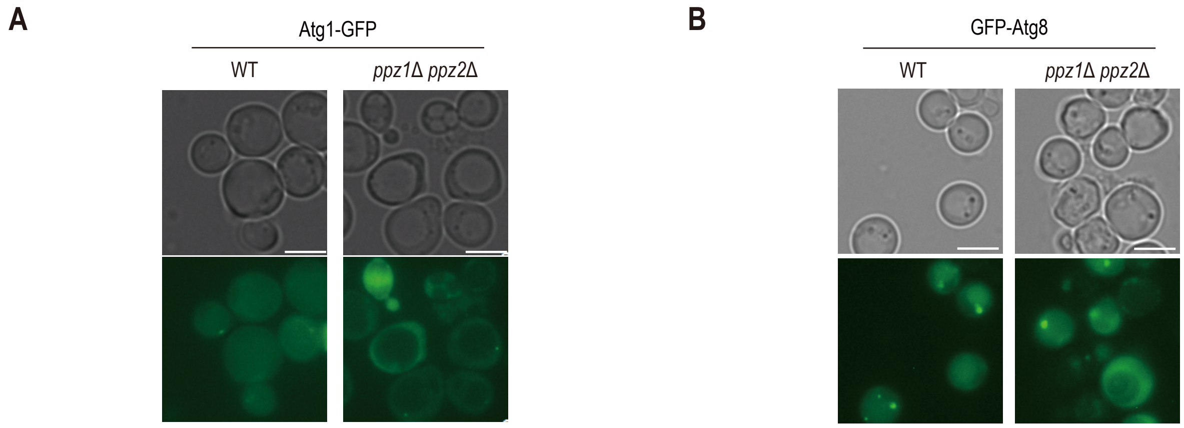

Supplymental Fig3(Supp of Fig. 3)

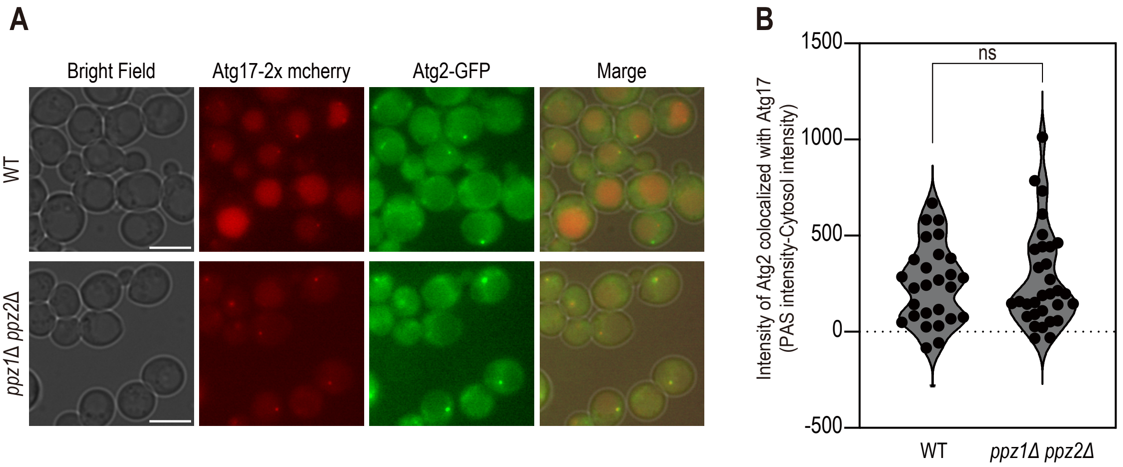

Supplymental Fig4(Supp of Fig. 4)

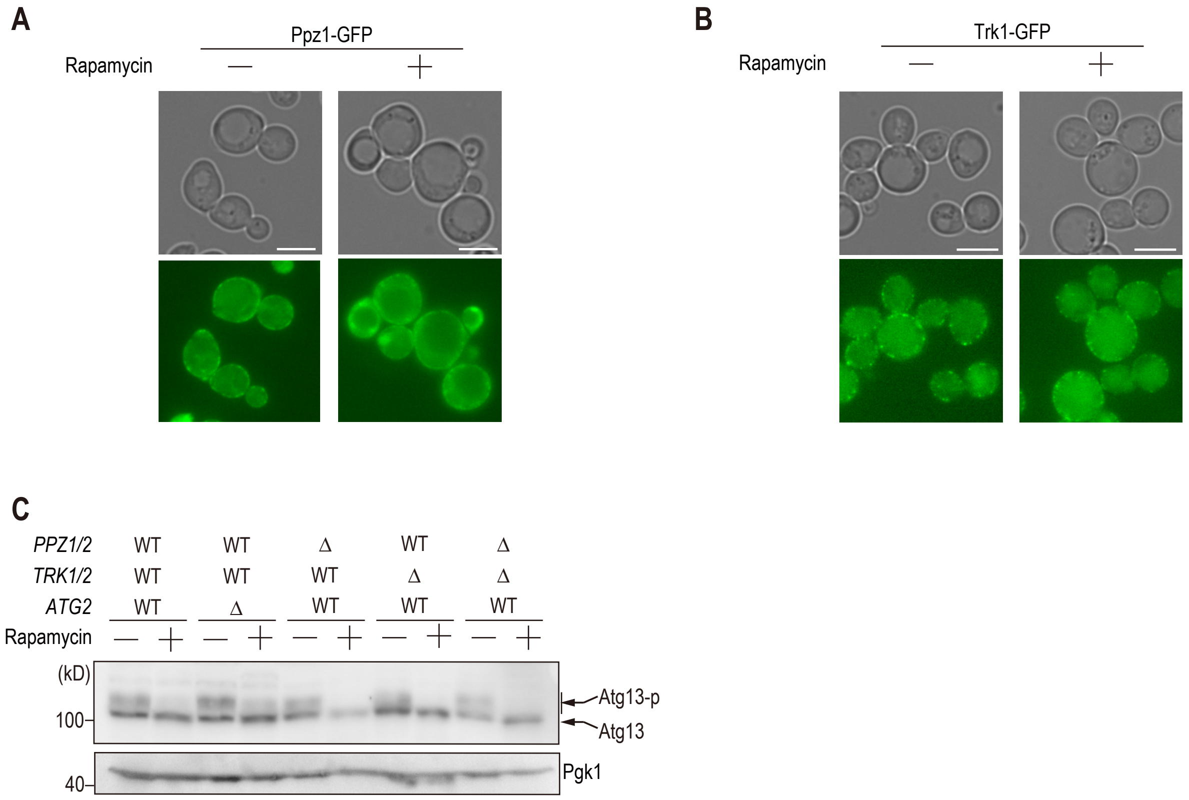

Supplymental Fig5(Supp of Fig. 5)

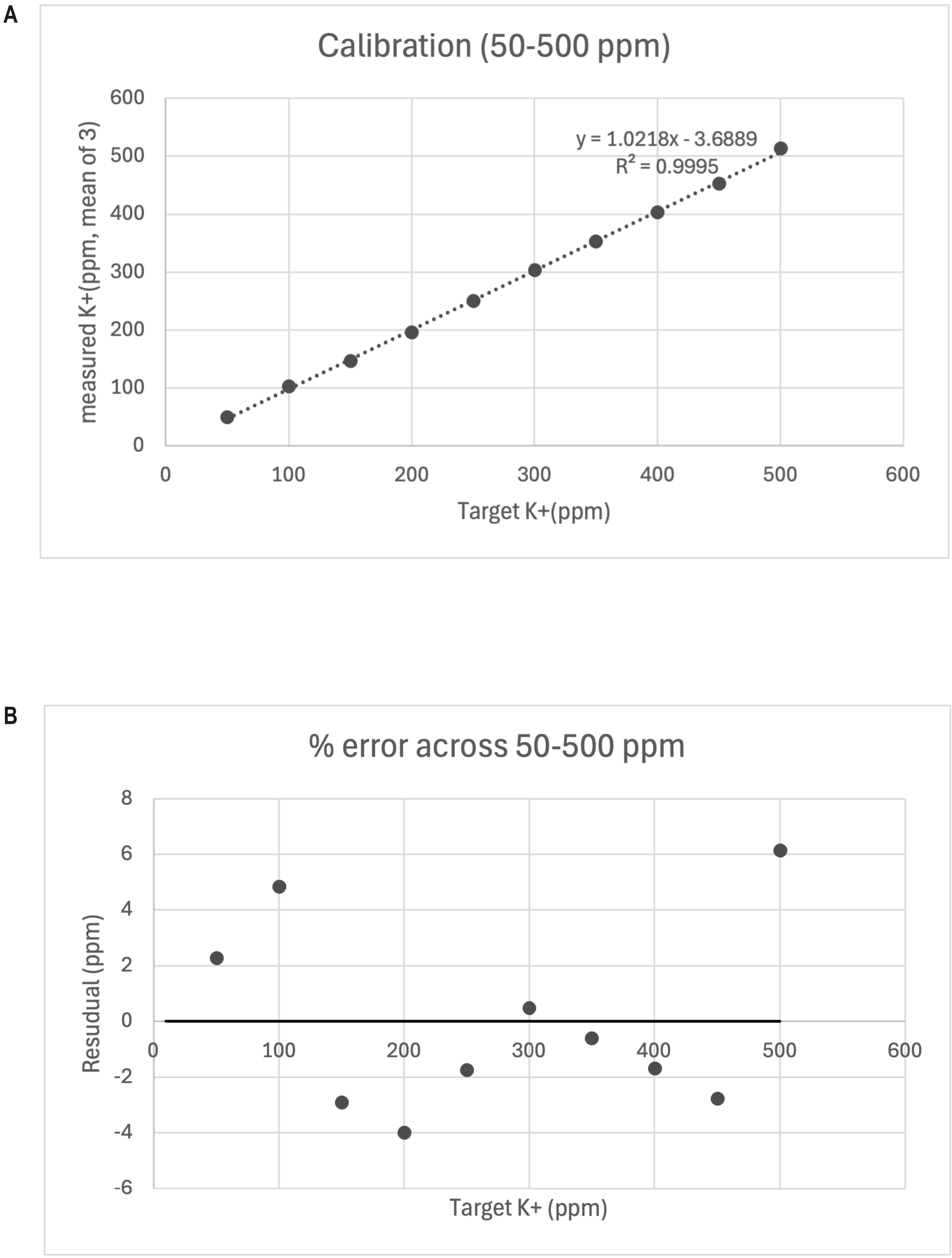
